# Environmental selection and spatiotemporal structure of a major group of soil protists (Rhizaria: Cercozoa) in a temperate grassland

**DOI:** 10.1101/531970

**Authors:** Anna Maria Fiore-Donno, Tim Richter-Heitmann, Florine Degrune, Kenneth Dumack, Kathleen M. Regan, Sven Mahran, Runa S. Boeddinghaus, Matthias C. Rillig, Michael W. Friedrich, Ellen Kandeler, Michael Bonkowski

## Abstract

Soil protists are increasingly appreciated as essential components of soil foodwebs; however, there is a dearth of information on the factors structuring their communities. Here we investigate the importance of different biotic and abiotic factors as key drivers of spatial and seasonal distribution of protistan communities. We conducted an intensive survey of a 10m^2^ grassland plot in Germany, focusing on a major group of protists, the Cercozoa. From 177 soil samples, collected from April to November, we obtained 694 Operational Taxonomy Units representing >6 million Illumina reads. All major cercozoan groups were present, dominated by the small flagellates of the Glissomonadida. We found evidence of environmental filtering structuring the cercozoan communities both spatially and seasonally. Spatial analyses indicated that communities were correlated within a range of four meters. Seasonal variations of bactevirores and bacteria, and that of omnivores after a time-lapse, suggested a dynamic prey-predator succession. The most influential edaphic properties were moisture and clay content, which differentially affected each functional group. Our study is based on an intense sampling of protists at a small scale, thus providing a detailed description of the niches occupied by different taxa/functional groups and the ecological processes involved.

## 1. Introduction

Our understanding of soil ecosystem functioning relies on a clear image of the drivers of the diverse interactions occurring among plants and the components of the soil microbiome - bacteria, fungi and protists. Protists are increasingly appreciated as important components of soil foodwebs (Bonkowski *et al.*, 2019). Their varied and taxon-specific feeding habits differentially shape the communities of bacteria, fungi, algae, small animals and other protists (Geisen 2016; Trap *et al.*, 2016). However, soil protistology is presently less advanced than its bacterial or fungal counterpart, and there is a dearth of information on the factors structuring protistan communities: this may be due to the polyphyly of protists, an immensely heterogeneous assemblage of distantly related unicellular organisms, featuring a vast array of functional traits (Pawlowski *et al.*, 2012).

Assessing how microbial diversity contributes to ecosystem functioning requires the identification of the appropriate spatial scales at which biogeographies can be predicted and the roles of homogenizing or selective biotic and abiotic processes. Local contemporary habitat conditions appear to be the most important deterministic factors shaping bacterial biogeographies, although assembly mechanisms might differ between different taxonomic or functional groups (Karimi *et al.*, 2018; Lindström and Langenheder 2012). Spatial distribution of microorganisms is driven by different factors at different scales (Meyer *et al.*, 2018). At a macroecological scale, soil microbial communities are mostly shaped by abiotic factors. Bacteria and archaea, in particular, are mostly influenced by pH (Kaiser *et al.*, 2016; Karimi *et al.*, 2018; Meyer *et al.*, 2018), while moisture and nutrient availability seems to drive the diversity and composition of fungal communities (Peay *et al.*, 2016). At a smaller scale, soil moisture can play a role for bacteria (Brockett *et al.*, 2012; Serna-Chavez *et al.*, 2013), while historical contingency and competitive interactions seem to be shaping fungal communities (Peay *et al.*, 2016). Seasonal variability is a major factor driving prokaryotic communities (Lauber *et al.*, 2013), and this has been confirmed at our study site for bacteria and archaea (Regan *et al.*, 2014; Regan *et al.*, 2017; Stempfhuber *et al.*, 2016).

However, due to high microbial turnover rates as well as their high capacity of passive dispersal, a great proportion of variation in microbial community assembly can be ascribed to stochastic processes and not deterministic ones (Nemergut *et al.*, 2013), although it is unclear to which degree (Dini-Andreote *et al.*, 2015). Despite climatic conditions regulating annual soil moisture availability have been shown to influence protistan communities at large scales (Bates *et al.*, 2013), some authors suggested that their assembly could be entirely driven by stochastic processes (Bahram *et al.*, 2016; Zinger *et al.*, 2018). Providing a thorough sampling of protistan communities in soil, we hypothesized that spatial and temporal abiotic and biotic processes would significantly contribute to protist community assembly.

Our study site was located in the Swabian Alps, a limestone middle mountain range in southwest Germany and part of a larger interdisciplinary project of the German Biodiversity Exploratories (Fischer et al., 2010). We applied a MiSeq Illumina sequencing protocol using barcoded primers amplifying a c. 350bp fragment of the hypervariable region V4 of the small subunit ribosomal RNA gene (SSU or 18S) (Fiore-Donno *et al.*, 2018). We focused on Cercozoa (Cavalier-Smith 1998) as an example of a major protistan lineage in soil (Domonell *et al.*, 2013; Geisen *et al.*, 2015; Grossmann *et al.*, 2016; Turner *et al.*, 2013; Urich *et al.*, 2008). This highly diverse phylum (c. 600 described species, (Pawlowski *et al.*, 2012)) comprises a vast array of functional traits in morphologies, nutrition and locomotion modes, and thus can represent the diversity of soil protists. We provided a functional trait-based classification of the cercozoan taxa found in our survey. We explored in a small, unfertilized grassland plot, how spatial distance, season, and edaphic parameters (abiotic and biotic) interact to shape the diversity and dynamics of the cercozoan communities.

## 2. Materials and Methods

### 2.1 Study site, soil sampling and DNA extraction

The sampling site was located near the village of Wittlingen, Baden-Württemberg, in the Swabian Alb (“Schwäbische Alb”), a limestone middle mountain range in southwest Germany. Details of the sampling procedure are provided elsewhere (Regan *et al.*, 2014; Stempfhuber *et al.*, 2016). Briefly, a total of 360 samples were collected over a 6-month period from spring to late autumn in a 10 m^2^ grassland plot in the site AEG31 of the Biodiversity Exploratory Alb (48.42 N; 9.5 E), with a minimum distance of 0.45 m between two adjacent samples (Fig. S1). For this study, we selected 180 samples, 30 samples from each sampling date (April, May, June, August, October, and November 2011). We were provided the soil DNA extracted from c. 600 mg/sample as described (Regan *et al.*, 2017). Soil physicochemical parameters were determined as described (Regan *et al.*, 2014), and the parameters included in our analyses are listed in Table S1, with their seasonal variation illustrated in Fig. S2. Over this area, spatial variability was limited; only the proportion of clay content varied (indicated in Fig. S2 by high boxes). Soil moisture changed most dramatically over the sampling period, with a peak in April and lowest values in May and October (average=40%, max=63%, min=23%, SD=11). Microbial biomass-related carbon and nitrogen parameters peaked in April. Bacterial cell counts showed a distinct peak in April, which was only partially reflected in the bacterial abundance as determined by qPCR. Living plant and fungi-related parameters followed the seasonal pattern of a minimum after winter, a maximum in summer, and a decrease in autumn (for the plant biomass, after mowing in August), with plant litter biomass following an inverse trend (Fig. S2).

### 2.2 Amplification, library preparation and sequencing

Primer design, barcoding primers, amplification, library preparation, and Illumina sequencing have been described in detail (Fiore-Donno *et al.*, 2018). The primers covered nearly the total diversity of Cercozoa, although they were biased against the parasitic Phytomyxea. The primers amplified a fragment of 335-544bp of the hypervariable region V4 of the SSU. Briefly, amplicons were obtained in two successive PCRs, the first using 1 µl of 1:10 soil DNA as template, the second using semi-nested primers and 1 µl of the first PCR as template. Barcoded primers were used in the second PCR to index samples. We pooled 15 μl of each of the successfully amplified samples (including the mock community, see below), then reduced the total volume to 80 μl; c. 800 ng of amplicons were used for the single library preparation as previously described (Fiore-Donno *et al.*, 2018). The library concentration was 23 nM, and 10 pM were used for the Illumina sequencing run. Sequencing was performed with a MiSeq v2 Reagent kit of 500 cycles on a MiSeq Desktop Sequencer (Illumina Inc., San Diego, CA, USA) at the University of Geneva (Switzerland), Department of Genetics and Evolution. To validate the bioinformatics pipeline (see below), we amplified DNA from a “mock community”, consisting of known representative Cercozoa (Fiore-Donno *et al.*, 2018), plus *Cercomonas longicauda* provided by S. Flues (GenBank DQ442884), totaling 11 species.

### 2.3 Sequence processing

Paired reads were assembled following a published protocol (Lejzerowicz *et al.*, 2014). The quality filtering discarded (i) all sequences with a mean Phred quality score < 30, shorter < 25 bases, with 1 or more ambiguities in the tag or in the sequence and with more than one ambiguity in the primers, and (ii) assembled sequences with a contig of < 100bp and more than 10 mismatches (Table 1). Sequences were sorted by samples (“demultiplexing”) via detection of the barcodes (Table S2) (Fiore-Donno *et al.*, 2018). The bioinformatics pipeline was optimized using the mock community. We first separated the sequences obtained from the 11 known taxa and ran the analysis with the steps listed in Table 1. The settings that made it possible to retrieve the expected 11 operational taxonomic units (thereafter OTUs) from the mock community were then applied to the environmental sequences.

**Table 1.**
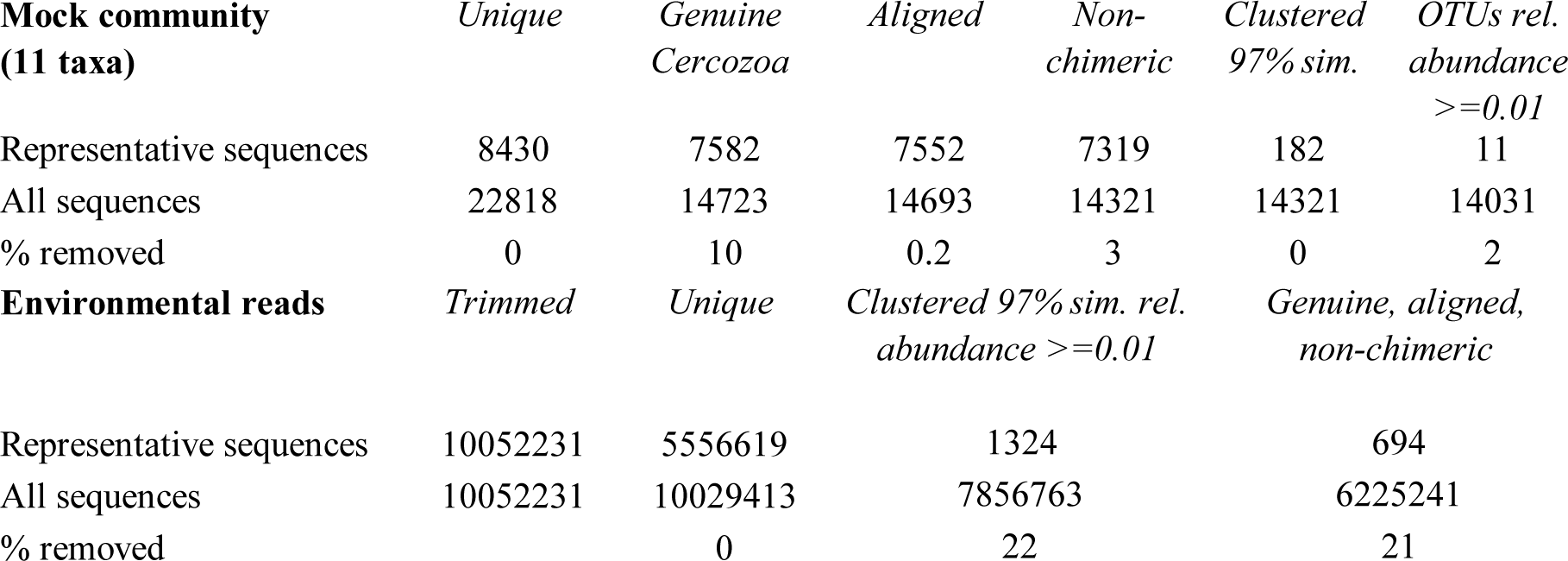
Number of reads retrieved at each step of the bioinformatic pipeline. Initial number of paired reads=15445807.

Sequences were clustered into OTUs using vsearch v.1 (Rognes et al., 2016), with the abundance-based greedy clustering algorithm (agc) implemented in mothur v.3.9 (Schloss et al., 2009) with a similarity threshold of 97%. Using BLAST+ (Camacho et al., 2008) with an e-value of 1^-50^ and keeping only the best hit, OTUs were identified using the PR2 database (Guillou et al., 2013); non-cercozoan OTUs were removed. Chimeras were identified using UCHIME (Edgar et al., 2011) as implemented in mothur v.3.9 as previously described (Fiore-Donno et al., 2018); chimeras and misaligned sequences were removed (Table 1).

### 2.4 Cercozoan functional traits

We selected three categories of ecological relevance: feeding mode, morphology, and locomotion mode. For the feeding mode, we classified the cercozoan OTUs into bacterivores, eukaryvores (feeding on algae, fungi, other protists and small animals but with no reports of feeding on bacteria) and omnivores, feeding on both bacteria and eukaryotes, according to information in the available literature. We applied two criteria for morphological classification: (i) the presence or absence of a shell (testate or naked); (ii) if the cell was an amoeba, a flagellate or an amoeboflagellate. We retained existing combinations, consisting of five categories (Table S3). We further distinguished two types of tests, organic or agglutinated - from those made of silica. The major difference in locomotion mode was set between cells creeping/gliding on substrate (on soil particles or on roots) versus free-swimming ones (in interstices filled with water). The phytomyxean parasites, due to their peculiar life cycle, were considered separately in each functional category. We assigned traits at different taxonomic levels, from order to genus (Table S3), and provide the respective references (Table S4).

### 2.5 Statistical and phylogenetic analyses

All statistical analyses were carried out within the R environment (R v. 3.5.1) (R Development Core Team 2014). Community analyses were performed with the packages vegan (Oksanen *et al.*, 2013) and betapart (Baselga 2017). We used: (i) Mantel tests and correlograms for spatial analysis; (ii) redundancy analysis (RDA) and generalized mixed effect models to describe the effect of environmental factors on community and population level; (iii) analysis of similarity (anosim), multi-response permutation procedure (MRPP) and multivariate permutational analysis of variance to analyze temporal variation at the community level; (iiii) post-hoc analysis (estimated marginal means on generalized square models with correction for spatial autocorrelation to test temporal variation of taxonomic and functional groups; (iiiii) variance partitioning to disentangle community turnover into spatial, temporal, and environmental components. Cercozoan diversity was illustrated using the Sankey diagram generator V1.2 (http://sankey-diagram-generator.acquireprocure.com/, last accessed Oct. 2018). Analyses are described in detail in Supplementary Data 1.

## 3. Results

### 3.1 Sequencing results

We obtained over 15 million filtered, paired reads (Table 1). The rate of mistagging during the sequencing run (indicating cross-over between adjacent clusters) was low, only 0.92%. To retrieve the 11 OTUs from the mock community, OTUs represented by ≤ 0.01% of the total sequences had to be removed; thus, this cutoff was applied to the environmental sequences. Not applying this cutoff would have drastically inflated the number of OTUs retrieved from the mock community (182 OTUs instead of 11). The percentage of non-cercozoan OTUs accounted for only 7.63% of the total sequences, confirming the high specificity of the primers. We obtained 694 genuine cercozoan OTUs from 177 grassland soil samples representing 6225241 sequences (Table 1). Three samples could not be amplified (Fig. S1). An NMDS analysis indicated that a sample was an outlier, thus it was omitted from subsequent analyses. The average number of OTUs per sample was 637 (maximum 681, minimum 407, SE 38.2). The average of sequences/soil sample was 35171 (maximum 153794, minimum 9808, SE 18.415), leading to an estimation of the coexistence of an average of c. 1000 cercozoan OTUs per gram of soil. We provide a database with the OTU abundance in each sample, their taxonomic assignment and their estimated functional traits (Table S3 and Table S4). The phylogenetic tree (Fig. S5) obtained with 176 reference sequences and 694 OTUs was rooted with Phytomyxea (Endomyxa); the vampyrellids (Proteomyxidea) and the Novel Clades 10-11-12, were paraphyletic to the monophyletic Cercozoa (93%). In Cercozoa, the main clades were recovered, although with low support. We were able to recover environmental sequences from nearly every clade of the tree.

### 3.2 Diversity of Cercozoa

At a high taxonomic level, the majority of the 694 OTUs could be assigned to the phylum Cercozoa (91% of the sequences) (Fig. 1), the remaining to Endomyxa (9%) and to the *incertae sedis* Novel clade 10 (Tremulida, 1%). Only 39% of the OTUs had 97-100% similarity to any known sequence in the PR2 database (Fig. S3). The 12 most abundant OTUs (>10000 sequences) accounted for 45% of the total sequences, while many low-abundant OTUs (243 < 1000 sequences) contributed to only 3% of the total sequences. These 12 OTUs were attributed to five orders: Glissomonadida (mostly Sandonidae), Cryomonadida (mostly two different *Rhogostoma* lineages), Plasmodiophorida (mostly *Polymyxa graminis*), Cercomonadida (*Eocercomonas* and *Paracercomonas*), and Spongomonadida.

**Figure 1.**
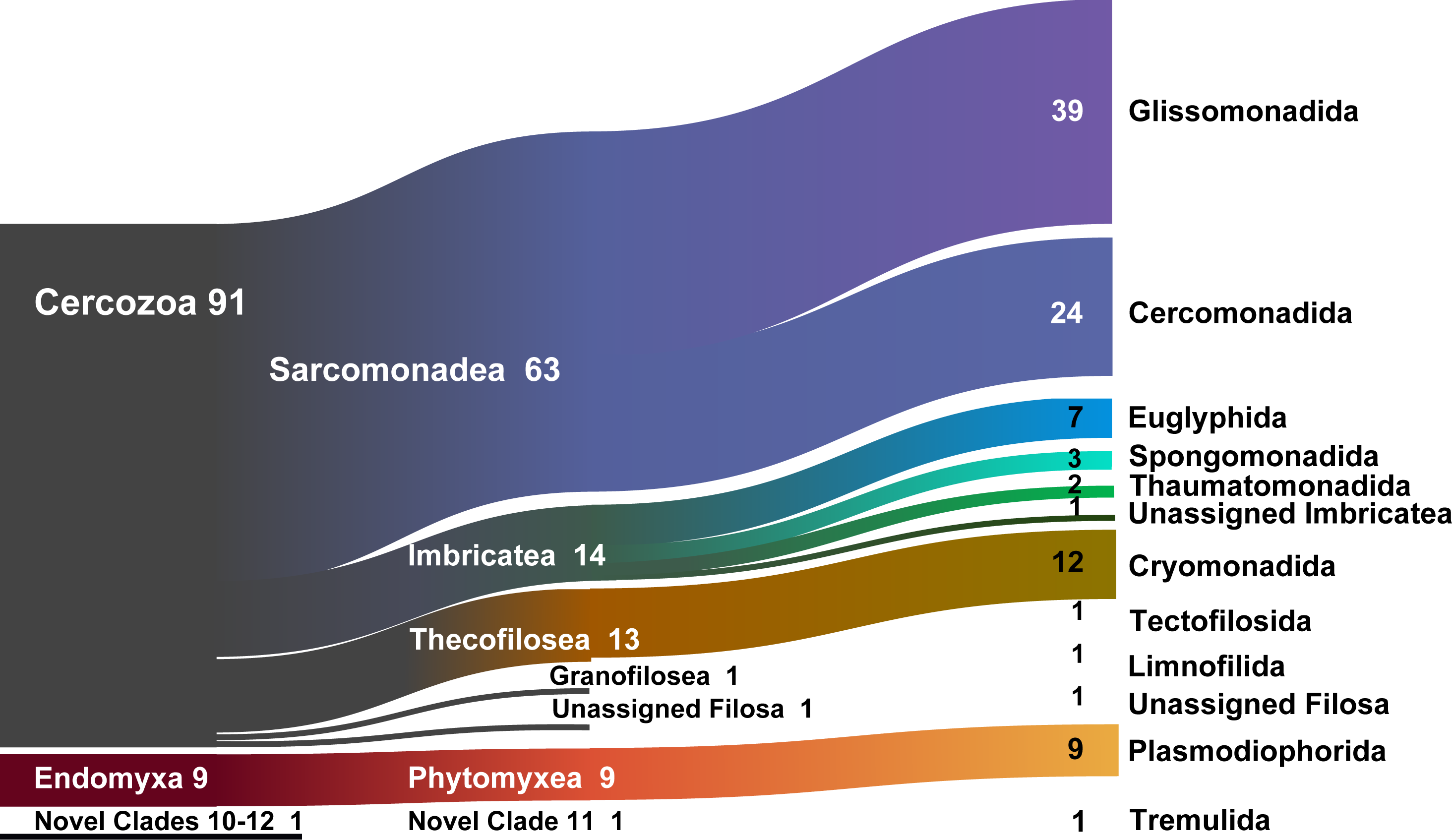
Sankey diagram showing the relative contribution of the OTUs to the taxonomic diversity. Taxonomical assignment is based on the best hit by BLAST. From left to right, names refer to subphylum (Filosa, Endomyxa), class (ending -ea) and orders (ending -ida). “unassigned” refer to sequences that could not be assigned to the next lower-ranking taxon or to “incertae sedis” order or families. Numbers are percentages of sequences abundances - taxa represented by <1% are not shown.

The rarefaction curve including all samples reached a plateau, suggesting that the global richness of 694 OTUs could have been obtained by already c. 70000 sequences (Fig. S4A), and by only 15 samples (Fig. S4B), and that the observed distribution patterns would not have been influenced by undersampling. Most OTUs were present in all sites (only 8.15 % of absences in Table S2). Multiple site beta diversity, calculated with either presence-absence or abundance on both rarefied and relative data, showed minor variation between the six sampling dates, which was confirmed by a resampling approach of random subsets (average resampled Bray-Curtis distances comprised between 0.715 and 0.745) (Table S5).

### 3.3 Functional diversity

More than half of the soil cercozoan OTUs were considered to be bacterivores (57%), followed by omnivores (21%), and eukaryvores (4%). Plant parasites (3%) and parasites of Oomycota (1%) were only marginally represented (Fig. 2). Naked flagellates (36%) or amoeboflagellates (34%) together constituted the majority of the morphotypes, whereas testate cells (organic/agglutinated or siliceous) were less frequent, 12 and 7% respectively. Naked amoebae were only marginally represented (1%). The dominating locomotion mode was creeping/gliding on substrate (86%), with only 1 % free-swimming.

**Figure 2.**
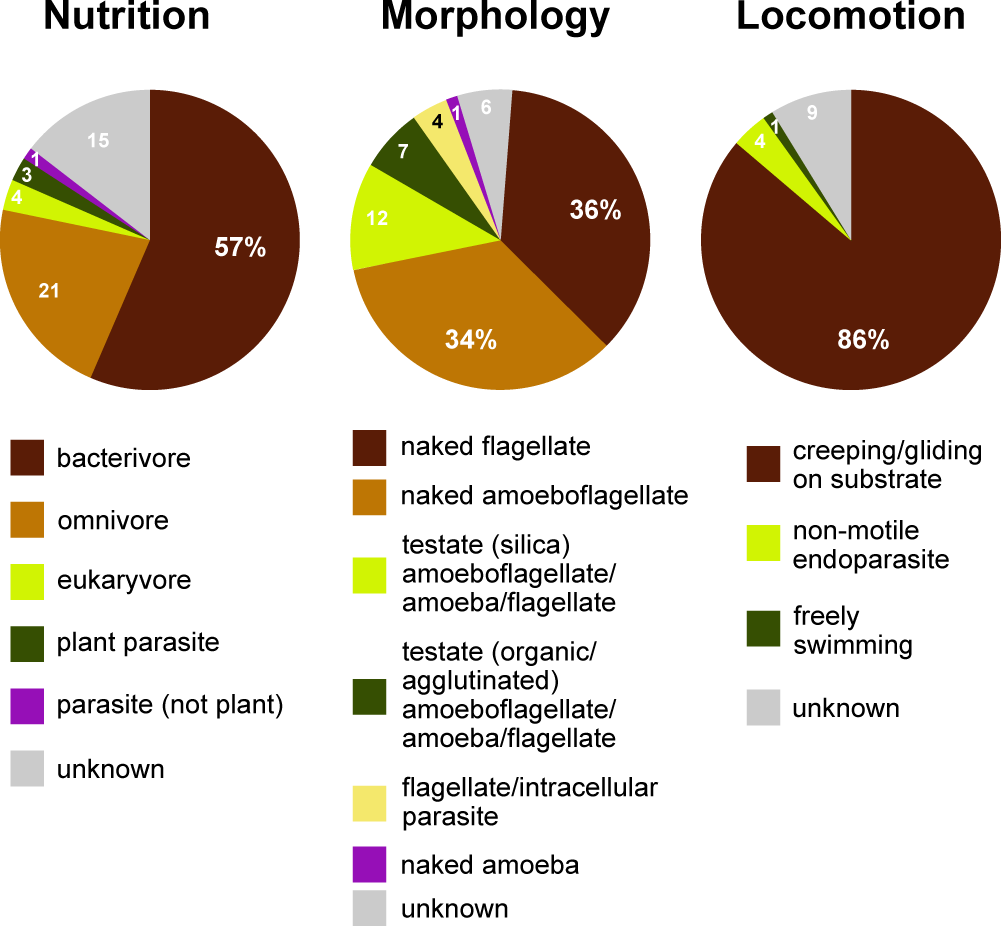
Functional diversity of cercozoan taxa. The relative proportions of functional groups classified according to nutrition, morphology and locomotion modes.

### 3.4 Spatial structuring and seasonal variation

The spatial distribution of the cercozoan communities was not random (Mantel test, r=0.05 to 0.18, p < 0.004), with the exception of flagellates. Similarity among communities decreased with distance, although the coefficients were low (Mantel correlograms, Fig. 3A). At distances between 0.45 to 3.9 m, cercozoan communities showed positive autocorrelations (see Table 2 for distance cutoffs), whereas communities at distances between 4.1 and 6.8 m were more dissimilar (i.e. negatively correlated). No spatial autocorrelations were observed at distances ranging from 6.9 to 12.4 m (Table 2). Similar results were obtained when functional traits were considered. The spatial structure of cercozoan communities differed slightly between seasons, with highest spatial variation in October and lowest in April and November (i.e. highest vs. lowest Mantel coefficients, Fig. 3B). Despite seasonal variation, the distance at which communities converged towards similarity corresponded to the smallest sampling distance (45 cm) for all seasons and with an increasing number of distance classes.

**Table 2.** Summary of Mantel correlogram (with ten distance categories) applied to the OTUs, comparing the effect of spatial distances, showing positive correlations up to a distance of 3.9 m. Significant correlations are in bold.

| Distance<br>classes (m) | All OTUs |  | gliding cells |  | testate cells |  |
| --- | --- | --- | --- | --- | --- | --- |
|  | Mantel<br>correlation r | p<br>(corrected) | Mantel<br>correlation r | p<br>(corrected) | Mantel<br>correlation r | p<br>(corrected) |
| 0.5 - 1.59 | 0.0656 | **0.0002** | 0.065 | **0.0001** | 0.069 | **0.0001** |
| 1.6 - 2.7 | 0.042 | **0.001** | 0.044 | **0.0005** | 0.047 | **0.0003** |
| 2.8 - 3.9 | 0.0421 | **0.0018** | 0.041 | **0.0022** | 0.044 | **0.0008** |
| 4.0 - 5.0 | -0.0483 | **0.001** | -0.05 | **0.0012** | -0.058 | **0.0004** |
| 5.1 - 6.2 | -0.0659 | **0.0002** | -0.066 | **0.0005** | -0.072 | **0.0005** |
| 6.3 - 7.4 | -0.0213 | 0.0642 | -0.021 | 0.0696 | -0.036 | **0.005** |
| 7.5 - 8.6 | -0.0735 | 0.3181 | -0.007 | 0.313 | -0.003 | 0.4309 |
| 8.7 - 9.7 | 0.0102 | 0.2893 | 0.012 | 0.5128 | 0.034 | 0.558 |
| 9.8 - 10.9 | 0.01 | 0.3069 | 0.014 | 0.6495 | 0.026 | 0.1354 |
| 11.0 - 12.4 | 0.005 | 0.3789 | 0.008 | 0.866 | 0.016 | 0.2332 |

**Figure 3.**
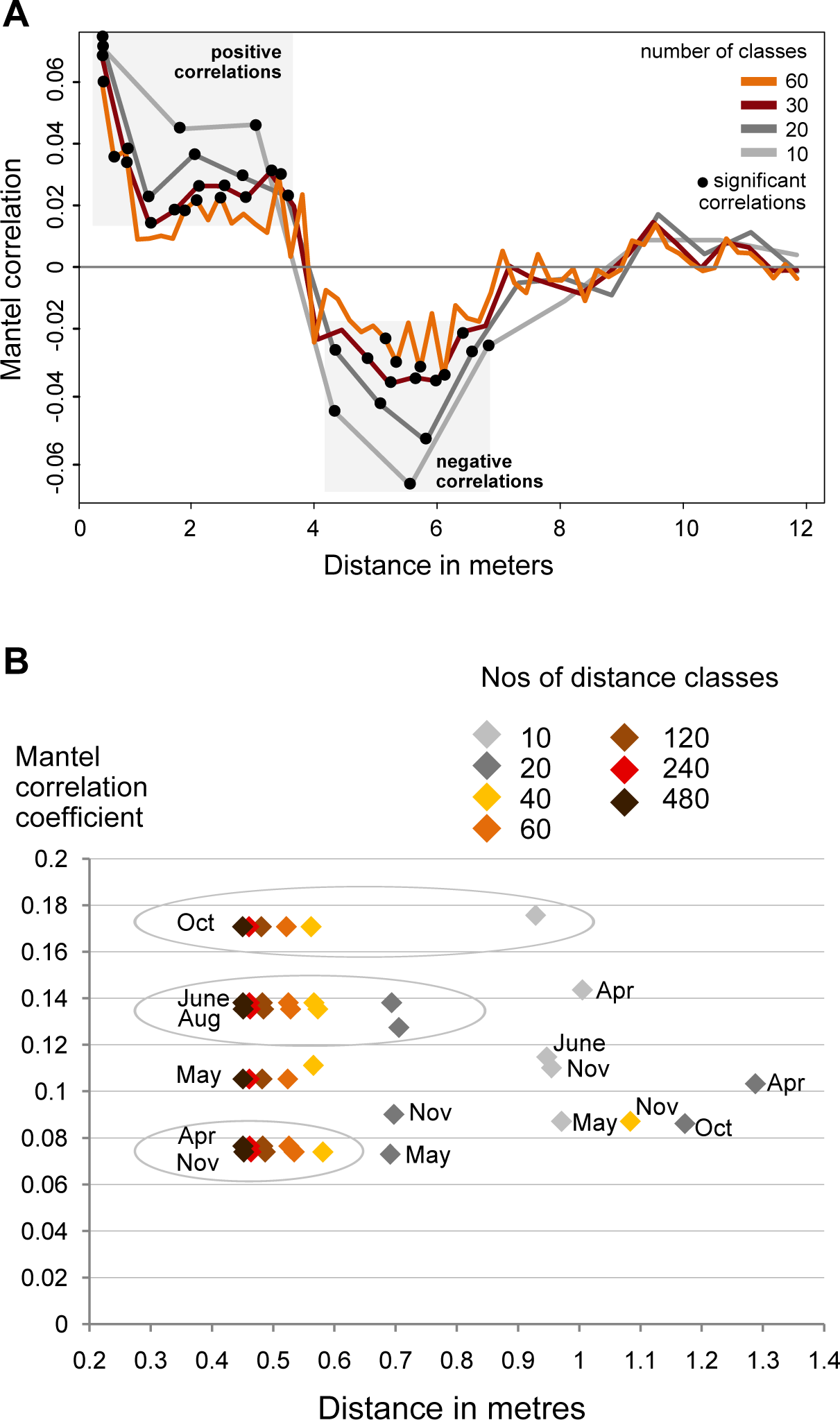
**A.** Mantel correlograms based on Bray-Curtis OTUs dissimilarities compared to Euclidian spatial distances and the spearman correlation coefficient showing how the similarities between cercozoan communities varied with distance. Only significant values (p < 0.05) are highlighted with black disks. Positive correlations occur at distances of 0.5-3.9 m, negative correlations at distances between 4.0-6.8 m. From 7 to 12.4 m, the communities varied randomly. **B.** Mantel correlograms as above, calculated for each season. Squares indicate the largest distance at which positive correlation were found. The smaller the intervals, the smaller the distance at which positive correlations occurred, converging towards the minimum distance of 0.45 m.

In contrast to the overall homogeneity of beta diversity, the cercozoan community structure changed significantly over time (anosim: *R=*0.22, p < 0.001; PERMANOVA: F_5-170=_5.009, p < 0.001; mrpp: p < 0.001). The cercozoan communities, binned by families or functional groups, showed distinct seasonal patterns of relative abundance (Fig. 4). While the bacterivorous families peaked in April and decreased in May (except Spongomonadidae), the omnivorous, shell bearing Euglyphidae, Rhogostomidae, and Trinematidae showed an opposite pattern, increasing from April to May. The relative abundance of plant parasites (*Polymyxa* and *Spongospora*) did not differ over the sampling season.

**Figure 4.**
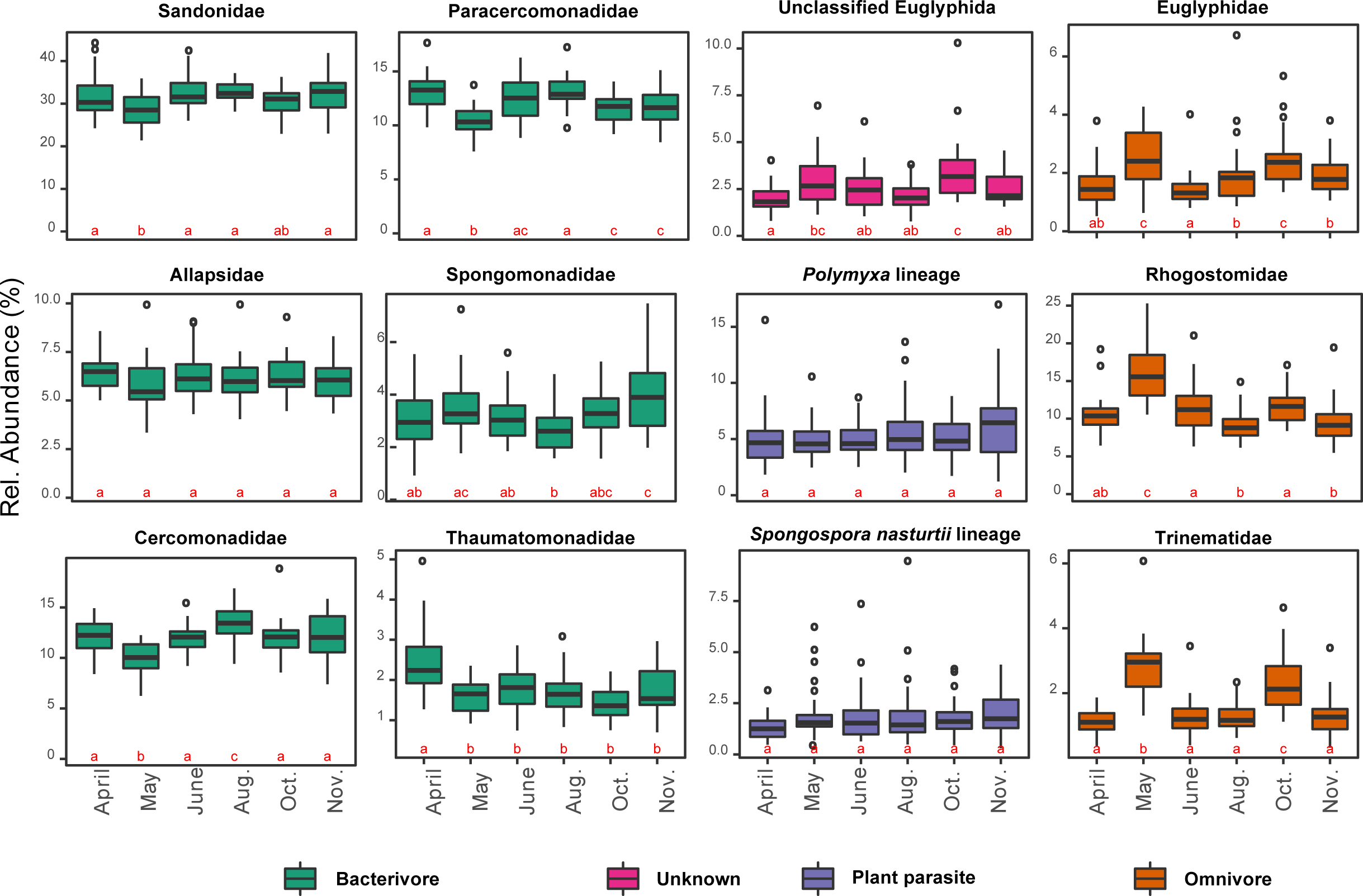
Box plots of the seasonal variation of the relative abundance of the 12 most abundant cercozoan families characterized by their nutrition mode (in different colors). Note that the y-scale varies between graphs. The small letters a, b and c indicate if the difference in abundance is statistically significant between months (contrasts of estimated marginal means on generalized least squares models after correction for spatial autocorrelation). A change of letter from “a” to “b” or “c” indicates a significant difference between months; “ab” indicate a non-significant difference, with values intermediate between “a” and “b”.

### 3.5 The main driver of cercozoan species turnover: season, distance or soil parameters?

Cercozoan beta diversity was influenced by soil parameters (RDA: F=4.162, p<0.001), spatial distance (RDA: F=2.202, p<0.001) and seasonality (RDA: F=3.389, p<0.001). Variance partitioning among the three predictors indicated that soil parameters, spatial distance and seasonality together accounted for 18% (adjusted R^2^) of the total variation in cercozoan beta diversity (Fig. 5). Soil abiotic factors (6%) and spatial distance (5%) explained roughly a similar proportion of the variation, followed by seasonality (2%).

**Figure 5.**
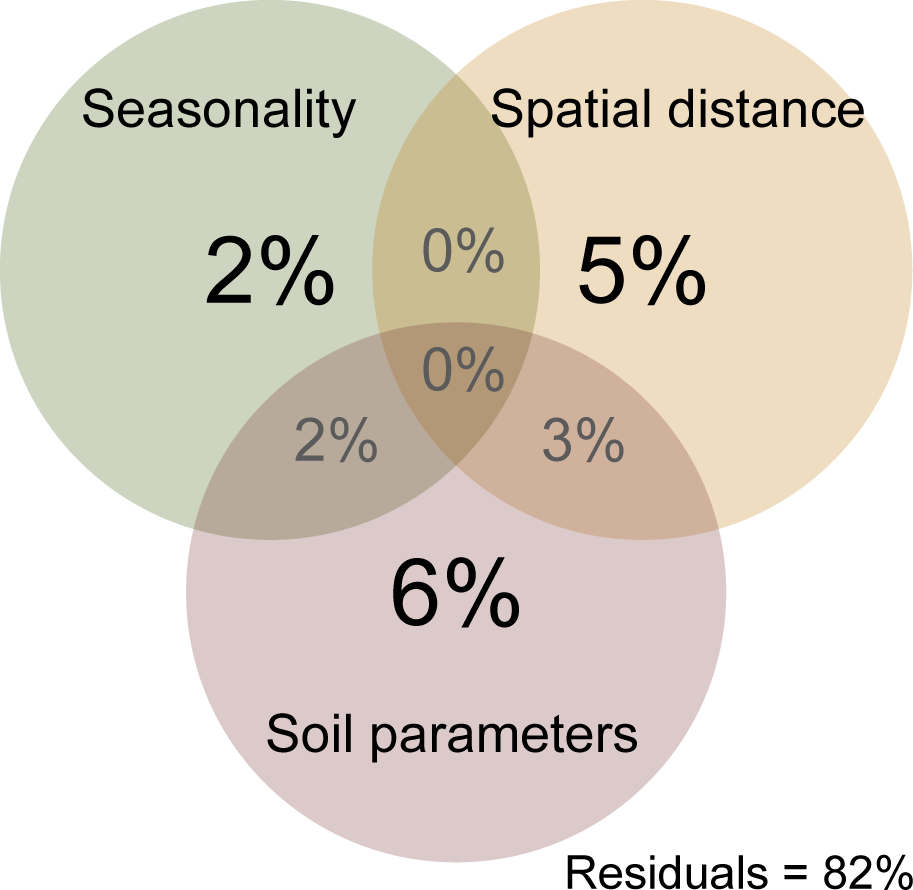
Venn diagram of variation partitioning analysis illustrating the effect of distance, season and environment (biotic and abiotic edaphic factors) on the cercozoan communities. Values show the percentage of explained variance by each set of variables, and of joined effects in the intersections.

### 3.6 Edaphic parameters influencing cercozoan communities

Among soil abiotic factors, soil moisture was most important (ANOVA: F_1,163_=11.75, p<0.001), followed by clay content, organic C, pH, litter, soil total N, and NO_3_ ^-^ content (Table 3). Biotic parameters (microbial biomass C and N contents, the abundance of archaeal and fungal PLFAs) explained a significant but lower amount of the variation in cercozoan beta diversity (Table 3). Using linear mixed effect models, we specified which abiotic and biotic factors affected the 12 most abundant cercozoan families (Table S6). Two-thirds of the cercozoan families (8 of 12) were positively affected by soil moisture (p<0.001), but not the Spongomonadidae, Allapsidae, and the parasitic *Spongospora nasturtii* and *Polymyxa* lineages. The relative abundances of the testate amoebae, i.e. Euglyphidae and unclassified Euglyphida, Trinematidae, and Rhogostomidae, responded negatively to soil moisture. In contrast, naked flagellates and amoeboflagellates were positively correlated with increasing soil moisture. Flagellates correlated negatively and amoeboflagellates positively with clay content. Flagellates, Sandonidae, and plant parasitic *Polymyxa* were positively correlated with pH, while testate amoebae (Euglyphidae) were negatively associated with pH (Table S6).

**Table 3.** ANOVA results of the most parsimonious RDA model including selected edaphic and biotic parameters.

| **Best predictor** | **F ratio<sub>1,163</sub>** | **p val** |
| --- | --- | --- |
| Soil moisture | 11.754 | <0.001 |
| % of clay | 4.927 | <0.001 |
| Soil organic C | 3.846 | <0.001 |
| pH | 3.385 | <0.001 |
| litter | 2.340 | <0.001 |
| Total N | 2.237 | <0.010 |
| NO<sub>3</sub><sup>-</sup> | 1.979 | <0.004 |
| N microbial biomass | 1.669 | <0.012 |
| Archaeal 16S<br>abundance | 1.566 | <0.023 |
| C microbial biomass | 1.430 | <0.043 |
| Fungal PLFAs | 1.405 | <0.053 |

## 4. Discussion

### 4.1 Environmental selection or random distribution?

Using Cercozoa as a model of very diverse and abundant soil protists, we demonstrated that their community composition in a small grassland site was nonrandom, but instead was spatially and temporally structured. With on average an estimated 1000 OTUs per gram of soil, high local species richness was a striking characteristic of cercozoan communities, with however small changes in beta diversity across space and time (Table S5). Our results tally with the consistent patterns of high alpha - and low beta diversity which have been found for protists in grasslands (Fiore-Donno *et al.*, 2016) and, including specifically Cercozoa, in tropical forests (Lentendu *et al.*, 2018). Sufficient sampling, attested by our saturation curves (Fig. S4), allowed us to further partition the beta diversity, proving the importance of deterministic processes of community assembly. Soil abiotic factors (6%) and spatial distance (5%) explained substantial variation indicating that cercozoan communities were significantly influenced by spatial gradients in the edaphic parameters (Fig. 5 & Fig. S2).

On a small spatial scale, our results correspond to large-scale patterns of protistan distribution as described by Lentendu et al. (2018), who established environmental selection as the main process driving protistan spatial patterns. This is in striking contrast to two recent studies. Bahram et al. (2016) reported a random spatial distribution of small soil eukaryotes in boreal forests, including protists, at a range of 0.01 to 64 m. They explained the observed distribution by invoking drift and homogenizing dispersal. The most striking difference compared with our study is the low efficiency of their ITS2 primers for the retrieval of protists (66 rhizarian OTUs, including the Cercozoa). This is at least one order of magnitude lower than in our data, possibly indicating a non-thorough sampling of the rhizarian communities in their study, which could in turn have hampered a robust assessment of the observed distribution patterns. In the second study, Zinger et al. (2018) suggested the absence of dispersal limitation and a stochastic distribution of protists in a tropical forest, using a 10 m-resolution sampling grid. This might have been too coarse to detect spatial patterns of protists, according to our results (assuming they apply to other environments), where no correlations could be detected at such a distance (Fig. 3).

Although we established the importance of environmental selection, homogenizing processes such as neutral assembly mechanisms may have contributed to the community assembly, as suggested by the positive autocorrelation of cercozoan communities up to a distance of four meters (Fig. 3). However, 18% of explained variance (Fig. 5) highlighted the importance of microhabitats for the nonrandom distribution of the cercozoan communities.

### 4.2 Habitat filtering and patch dynamic selected for specific functional traits

Significant relationships between functional traits and soil abiotic and biotic factors (Table S6) indicated trait selection by habitat filtering as an important driver of community composition in our study site. Some of these filters showed marked spatial gradients (clay content, pH, total N), while other showed more temporal variation (moisture, NH_4_^+^, total plant biomass) (Fig. S2). Soil moisture is well known to influence microbial activity (Tecon and Or 2017). The spatial variation in clay content together with the temporal variation in soil moisture triggered opposite responses from specific lineages or functional groups (Table S6 and Fig. 6). While flagellates and amoeboflagellates were favored by moisture, the cercozoan testate cells were correlated with drier conditions, suggesting patch dynamic processes connected to different living modes. In accordance with our model for Euglyphidae, Ehrmann et al. (2012) reported preference of testate amoebae for relatively dryer soils and lower pH in forests. We can conclude that in grasslands, testate amoebae exhibit drought resistance, in contrast with naked cells and with cells covered with scales (i.e. Thaumatomonadidae, suggesting a protective role of the shell (Table S4). Building a shell, however, slows down the reproduction rate (Schönborn 1992), and thus generates a trade-off between protection and reproductive fitness. In contrast, amoeboflagellates and flagellates, with their faster reproduction rate (Ekelund and Rønn 1994), would have a better fitness in moist conditions.

**Figure 6.**
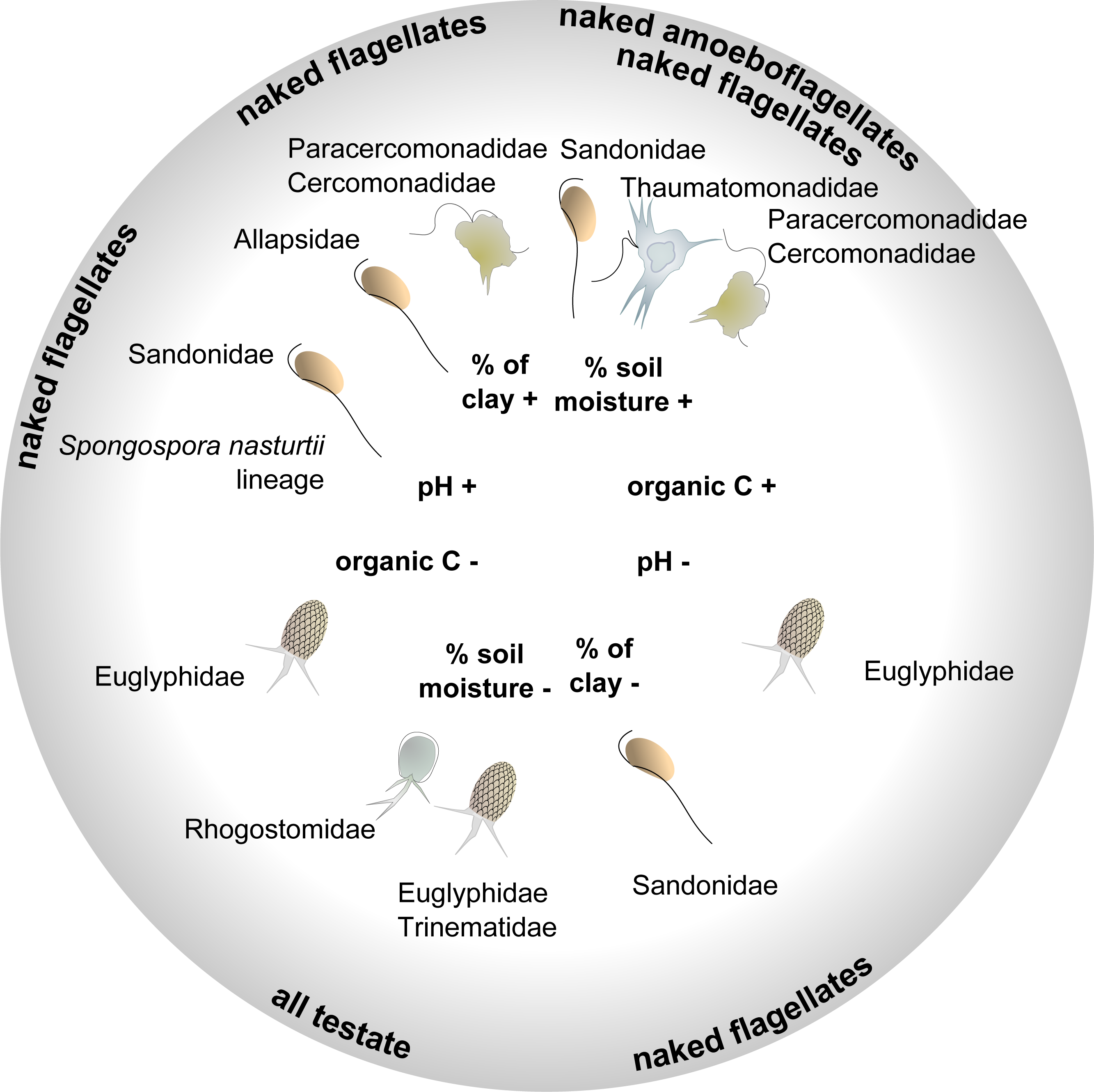
Schematic illustration showing the positive (+) or negative (-) interactions between the four most influent soil physicochemical parameters (based on the ANOVA, Table 3), and the relative abundance of the major families or morphotypes (details of the models in Table S6).

The spatial distribution of soil clay content was an important structuring factor in our study site. Experimentally increasing clay content has been shown to improve water retention capacity and favor soil bacteria (Heijnen *et al.*, 1993). In addition, it leads to a reduction of the habitable pore space, which in our study seemed to favor the amoeboflagellates (Paracercomonadidae and Cercomonadidae) and the naked flagellates (Allapsidae, but not Sandonidae) (Fig. 6 and Table S6). The small percentage of free-swimming protists was negatively correlated with bulk density (Table S6), thus favored by larger soil pores. Another important structuring factor was the pH, despite its little variation (Fig. S2 and Table S1 – pH 6.08-7.23). Globally, cercozoans have been shown to prefer neutral or basic soils (Dupont *et al.*, 2016) and their relative abundance was seen to increase significantly along a pH gradient from c. 4 to 6.5 (Shen *et al.*, 2014). In a reduced gradient, we showed that Sandonidae and the parasitic *Spongospora nasturtii* lineage were positively correlated with a slightly basic pH, and the Euglyphidae with a slightly more acidic one (Fig. 6 and Table S6). Our results thus suggest that different taxa could have different pH preferences. Soil organic carbon had a negative effect only on the Euglyphidae, in conjunction with nitrogen content, while the C/N ratio had a positive effect, suggesting their preference for low-nutrient soils with a relatively higher C than N content.

Bacterial cell counts were a major explanatory biotic factor. Bacterial numbers positively and predictably influenced the abundance of bacterivores but also that of eukaryvores (probably by an indirect effect). In our study, the majority of the cercozoan taxa that positively correlated with bacterial numbers are bacterial consumers, that were also identified as creeping/gliding on substrate. This is in accordance with soil protists feeding mostly on bacterial biofilms (Böhme *et al.*, 2009). Contrary to other bacterivores, the Spongomonadidae did not follow the seasonal variation of the bacteria (Fig. 4). This may be explained by their living modes, mostly in substrate-attached colonies (Strüder-Kypke and Hausmann 1998). It has been suggested that the colonial species may also feed by saprotrophy (Strüder-Kypke and Hausmann 1998), a hypothesis supported by the positive correlation with extractable organic carbon which we observed (Table S6). We know very little about the interactions of archaea with soil protists. Thus it is worth pointing out the so far unexplained significant negative effect of archaeal abundance on Paracercomonadidae (Table S6).

### 4.3 Seasonal variability affected the trophic structure of cercozoan communities

Cercozoan communities showed seasonal oscillations, in accordance with results from bacteria in grasslands (Muller Barboza *et al.*, 2018). Between April and May, a series of changes in edaphic factors was observed (Fig. S2). The high soil moisture in April favored bacterial activity and proliferation. At the same time, high levels of extractable organic C indicated abundant root exudates or decomposition at the onset of the plant growing season, when plant total biomass was still low. This likely triggered a series of events: the bacteria, usually C limited, were stimulated by this C input. Since fungi were not yet abundant, bacteria were mostly responsible for the high nitrogen content in the microbial biomass. Consequently, the release of NH_4_^+^ (also peaking in April) may be best explained by protistan predation on bacteria (Bonkowski 2004). This was confirmed by the high abundances of five bacterivorous families (Fig. 4). In May, bacteria, bacterivores, and nitrogen-related parameters all decreased, together with soil moisture. In sharp contrast, all three omnivorous families increased in May. We hypothesize that the predation of the omnivores could have contributed to the decline of the bacterivores, in addition to the negative effect of declining soil moisture (Fig. S2).

Our hypothesis that the abundance of plant parasites would follow the annual plant cycle was not supported, since they showed no seasonal variation (Fig. S2). Phytomyxeans are known to form resistant cysts in plant root hairs that remain for years in the soil after plant decay (Dixon 2009). Our study, based on DNA, does not make it possible to distinguish between active and resting stages, but we probably also amplified extracellular DNA from recently deceased cells. Thus, the seasonal variation of cercozoans we observed was probably an underestimate.

### 4.4 Number of OTUs, diversity and functional traits

Cercozoan diversity was in line with previous studies, which established Sarcomonadea (Glissomononadia and Cercomonadida) as the dominant class in different terrestrial habitats (Fiore-Donno *et al.*, 2018; Geisen *et al.*, 2015; Howe *et al.*, 2009), and more specifically in feces (Bass *et al.*, 2016), in the soil of neotropical forests (Lentendu *et al.*, 2018), on the leaves of Brassicaceae (Ploch *et al.*, 2016), in heathlands (Bugge Harder *et al.*, 2016), and in German grasslands, including the site studied here (Venter *et al.*, 2017). Especially the glissomonads (mostly) bacterivorous small flagellates, seem to dominate in all types of grasslands, where they can reach 5% of all protistan sequences (Geisen *et al.*, 2015). However, the dominance of the remaining taxa is more variable between habitats. The Sarcomonadea were followed by the (mostly) omnivorous amoeboflagellates in cercomonads and by Cryomonadida, composed of amoebae or amoeboflagellates with organic, transparent tests. The widespread presence of Cryomonadida in soil has been overlooked in observation-based inventories, but confirmed by molecular environmental sampling (Bass *et al.*, 2016; Bugge Harder *et al.*, 2016; Lentendu *et al.*, 2014; Ploch *et al.*, 2016); especially the genus *Rhogostoma*, which is very common in soil (Fiore-Donno *et al.*, 2018).

In conclusion, we showed that environmental selection driven by abiotic and biotic edaphic factors significantly determined community assembly of Cercozoa. Considering functional traits and their trade-offs, we could highlight the importance of habitat filtering and patch dynamics as underlying processes. We believe that our study has bearing for other soil protists and soil ecosystems beyond the limits of this small grassland plot. Once the patterns underlying the small-scale distribution of protists are detected, they can be upscaled and contribute to understanding global protistan biogeographies. This is a perequisite for predicting effects of human-induced changes (i.e. land management or global warming) on these widespread and functionally important soil organisms.

## 5. Data accessibility

Raw sequences have been deposited in Sequence Read Archive (SRA, NCBI) SRR6187016 and BioProject PRJNA414535; the 694 OTUs (representative sequences) under GenBank accession # MG242652-MG243345.

## Supporting information

Suppl.Data1andFigS1-S5

Suppl.TablesS1-S6 Except S3

Table S3 Database

## 6. Acknowledgments

We thank Franck Lejzerowicz for designing the barcodes for the primers. Sebastian Flues, Kenneth Dumack and Sebastian Hess for providing the DNAs of the mock community. At the University of Geneva (CH), we thank Jan Pawlowski, Emanuela Reo and Ewan Smith. At the University of Hohenheim, we thank Doreen Berner and Robert Kahle for soil sampling and technical support in the laboratory. We thank the manager of the Alb Exploratory, Swen Renner, and all former managers for their work in maintaining the plot and project infrastructure; Simone Pfeiffer for giving support through the central office, Jens Nieschulze and Michael Owonibi for managing the central data base, and Markus Fischer, Eduard Linsenmair, Dominik Hessenmüller, Daniel Prati, Ingo Schöning, François Buscot, Ernst-Detlef Schulze, Wolfgang W. Weisser and the late Elisabeth Kalko for their role in setting up the Biodiversity Exploratories project. Field work permits were issued by the responsible state environmental offices of Baden-Württemberg.

## 7. Author contributions

EK and SM developed the design of the SCALEMIC experiment. KMR performed DNA extractions. KMR and RSB analyzed soil properties. AMFD conducted the amplifications, Illumina sequencing and bioinformatics pipeline. TRH, FD and AMFD performed statistics. AMFD, TRH and MB interpreted the data. AMFD wrote the manuscript. All co-authors approved the final version of the manuscript.

## 9. Funding

This work was partly funded by the DFG Priority Program 1374 “Infrastructure-Biodiversity-Exploratories”. Funding to AMFD and MB was provided by BO 1907/18-1; funding to EK, SM and RSB was provided by KA 1590/8-2 and KA 1590/8-3; funding to FD and MR was provided by the BiodivERsA grant “Digging Deeper”. We are liable to the Swiss National Science Foundation Grant 316030 150817 for funding the MiSeq instrument at the University of Geneva (CH).

**Supplementary Information** accompanies this paper on The ISME Journal website (http://www.nature.com/ismej)

## Supplementary material

**Supplementary Data 1**. Detailed description of the statistical and phylogenetic analyses.

### Supplementary tables

**Table S1**. Environmental parameters from the study site as in (Regan *et al.*, 2014) used in our statistical analyses. Their seasonal variation is shown in Fig. S2.

**Table S2**. Combinations of barcodes used in this study, with the corresponding samples.

**Table S3**. Database of the abundance of each OTU per sample. The taxonomic assignment (supergroup, class, order, family, genus and species) is provided according to the best hit by BLAST (PR2 database), with the % of similarity. Functional traits (morphology, nutrition and locomotion modes) were estimated following Table S4.

**Table S4**. References for the functional traits of the cercozoan taxa identified in this study.

**Table S5**. Beta diversity indices calculated for each sampling date.

**Table S6**. Linear mixed models showing the effects of the environmental predictors on the most abundant 12 cercozoan families, the morphotype, nutrition and locomotion modes. We give: a) the spatial correlation structure best correcting the starting model according to the AIC; b) the number of models within two AICc units (after model dredging); c) the number of predictors included in all models extracted in a); d) the remaining, highly significant predictors after fitting a model with just the consensus predictors in b), and their effect type (positive or negative); e) their significance level (p values: *<0.05, **<0.01, ***<0.001).

### Supplementary figures

**Figure S1**. Sampling design of the 10m^2^ grassland study site (Fig. A2 in Regan *et al*., 2014), with the 360 samples collected at six different dates and the spatial coordinates used to build the distance matrix. For this study, we selected the samples from the 15 areas outlined in grey (12 samples/area=180 samples). The DNA from samples 125, 185 and 305 from area 3 could not be amplified.

**Figure S2**. Box plots showing the seasonal variation of the environmental parameters from Table S1. Values were normalized (mean=0, SD=1).

**Figure S3**. Similarities of the OTUs with known sequences. OTUs are classified according to their percentage of similarity to the next kin by BLAST. The horizontal bar length is proportional to the number of OTUs in each rank. Shaded area=OTUs with a similarity ≥ 97%.

**Figure S4**. Description of the diversity. **A**. Rarefaction curve describing the observed number of OTUs as a function of the sequencing effort; saturation was reached with c. 70000 sequences. **B**. Species accumulation curve describing the sampling effort; saturation was reached with 15 samples.

**Figure S5**. Cercozoa Maximum Likelihood phylogenetic tree, inferred from an alignment of 870 taxa and 1,447 positions. The tree is rooted between Phytomyxea and the remaining taxa. OTUs found in this study are in bold. Main clades are named according to the PR2 database taxonomy, clades for which no OTUs were found are named in gray. Bootstrap values are given for each node. The scale bar indicates the fraction of substitutions per site.

