## Supplementary material for "Environmental selection and spatiotemporal structure of a major group of soil protists (Rhizaria: Cercozoa) in a temperate grassland": Suppl.Data1andFigS1-S5

**Supplementary Data 1. Detailed description of the statistical and phylogenetic analyses**

All statistical analyses were carried out within the R environment (R v. 3.5.1) (R Development Core Team 2014). Analyses were performed with the package vegan (Oksanen *et al.,* 2013), if not stated otherwise.

To evaluate if more sampling and sequencing effort would have revealed more richness, we carried out a rarefaction analysis based on OTUs accumulation curves, function *specaccum*, method rarefaction and 1000 random permutations; species richness was extrapolated using the function *specpool*. Indices of alpha-diversity, i.e. species richness and linearized Simpson diversity (Hill numbers $^{0}D$ and $^{2}D$, respectively), were computed from the raw OTU table (Table S3) using the function *iNEXT* for each season, with default settings (50 bootstraps, confidence interval 0.95) at a common sample size of twice the smallest sample in the dataset (Chao *et al.,* 2014).

Beta diversity was calculated as multiple site dissimilarities for each sampling date and for functional groups with functions *beta.multi* (presence-absence data, Sorensen index); with *beta.multi.abund* (abundance data, Bray-Curtis dissimilarities) for the total dissimilarities weighted by abundance; and with *beta.sample.abund*, which resamples Bray-Curtis dissimilarities from random subsets of 20 sites per sampling date with 100 permutations (abundance data) - all functions from package betapart (Baselga 2017).

### 1.1.1 Spatial analyses

The effect of distance on beta diversity was assessed using a Mantel test with 5,000 permutations (function *mantel*), between the Bray-Curtis dissimilarities matrix described above and the Euclidian distance matrix based on sample coordinates (function *dist*, Figure S1). To identify the smallest distance at which spatial correlations were found, we calculated Mantel correlograms using the Spearman correlation coefficient (function *mantel.corelog*, 9999 permutations). We further divided the spatial distances into increasingly smaller units - from 10 to 480 categories, with, however, increasing numbers resulting in many void categories. To take into account the seasonal variation, we also calculated the correlograms for each sampling date.

### 1.1.2 Seasonal variation

To test whether seasonal variation affected beta diversity, we performed multiple tests using the normalized OTU table: analysis of similarity (anosim), multi-response permutation procedure (MRPP) and multivariate permutational analysis of variance (PERMANOVA), all carried out with 999 permutations (functions *anosim*, *mrpp*, *adonis2*, respectively). The response of different taxonomic groups to seasonal change was examined with estimated marginal means (EMM, function *emmeans*, package emmeans) (Lenth 2018) on general least square models (GLS, function *gls*, package nlme) (Pinheiro *et al.,* 2018). Correction for spatial autocorrelation was done by selecting the best match out of four different correlation structures (spherical, gaussian, rational, exponential) based on the Akaike Information Criterion (AIC). The best model, i.e. with the lowest AIC, was then used to obtain EMM-based contrasts between sampling dates.

### 1.1.3 Environmental variables

The influence of environmental variables on specific groups of cercozoans (normalized abundances) was assessed using general linear mixed models (GLMM) where sampling date was introduced as a mixed random effect (function *lme*, package nlme) to account for seasonal change. As for the GLS models described above, correction for spatial autocorrelation was done by selecting the best model based on the lowest AIC value. Variable selection was carried out as follows: the best environmental predictors were preselected using the forward and backward selection approach (function *stepAIC*, package MASS) (Venables and Ripley 2002). The set of candidate predictors was subsequently used for multi-model inference, based on the corrected AIC (AICc) (function *dredge*, package MuMIn) (Barton 2018). This procedure yielded a list of potential models ranked by AICc. From this list, we considered a variable as potentially important if it appeared in all models within an interval of two units of AICc from the best scoring model (function *model.avg*, package MuMIn). In a final step to obtain even more sparse models, these "consensus" variables were only selected if they were found to be highly significant ($p<0.01$) in a final GLMM call.

### 1.1.4 Variance partitioning

Variance partitioning among environmental parameters, spatial and temporal components was assessed using the function *varpart* and visualized in a Venn diagram. Prior to the analysis, the OTU table was Hellinger-transformed (function *decostand*) and environmental data were normalized. Spatial and temporal structures in the response data (OTUs) were previously detected using Principal Coordinates of Neighbor Matrices (PCNM) (Borcard and Legendre 2002; Borcard *et al.,* 2004; Dray *et al.,* 2006). Briefly, an Euclidean distance matrix was calculated from spatial and temporal data and truncated to obtain a new set of independent spatial and temporal PCNM variables (neighbor matrix). For each set of PCNM variables, a principal coordinate analysis (*cmdscale* function) was carried out to select those PCNMs with positive eigenvalues. The resulting sets of PCNM variables were used in a redundancy analysis (function *rda*) with the *nondetrended* response data. The significant spatial and temporal PCNM variables were selected using the *forward.sel* function and the resulting model was assessed using the *anova* function. Constrained analysis was performed similarly, exept that the significant environmental variables were selected using the *ordiR2step* function. The resulting parsimonious model was tested using the *anova* function and the significance of each parameter was determined using the anova function *by term*. Variance partitioning (Peres-Neto *et al.,* 2006) among the significant PCNM (spatial and temporal) and environmental variables was carried out using the function *varpart*.

### 1.1.5 Phylogenetic analyses

The 694 OTUs obtained were added to a published reference alignment of nearly complete 18S sequences of 176 cercozoan taxa (Fiore-Donno *et al.,* 2018). The alignment had 1447 positions, 1013 alignment patterns, and a proportion of undetermined characters of 66.28%. Maximum likelihood (ML) analyses were run using RAxML 8.2.4 (Stamatakis 2014) with the GTR model of substitution and a 25 rate category discrete gamma distribution. The best scoring ML tree was inferred from 200 randomized starting Maximum Parsimony trees using the GTRMIX model. The best-scoring tree was used to report the confidence values as percentages obtained through 500 non-parametric bootstraps.

**Figure S1. Sampling design of the 10m$^2$ grassland study site** (Fig. A2 in Regan et al. 2014), with the 360 samples collected at six different dates and the spatial coordinates used to build the distance matrix. For this study, we selected the samples from the 15 areas outlined in grey (12 samples/area = 180 samples).

The DNA from samples 125, 185 and 305 from area 3 could not be amplified.

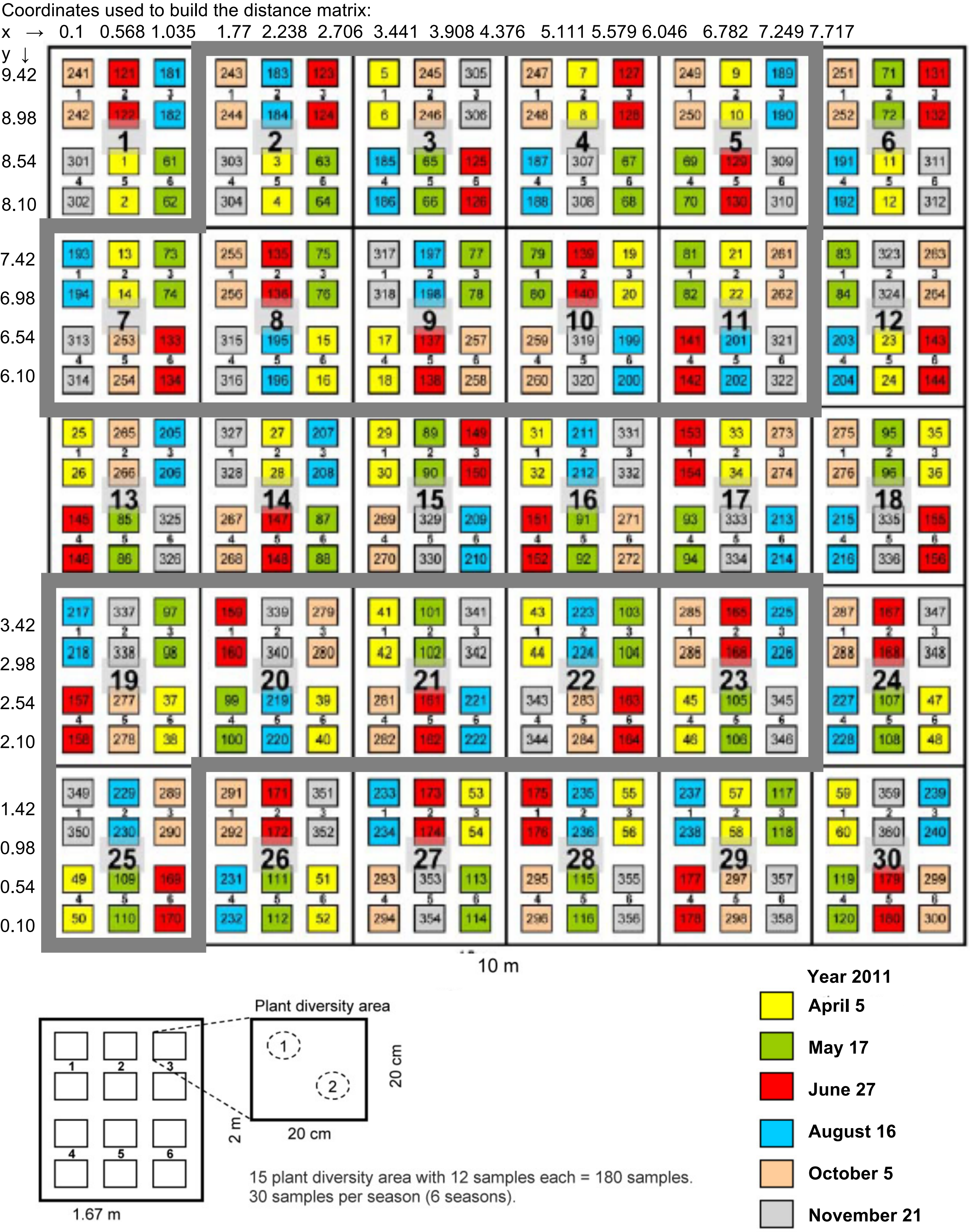

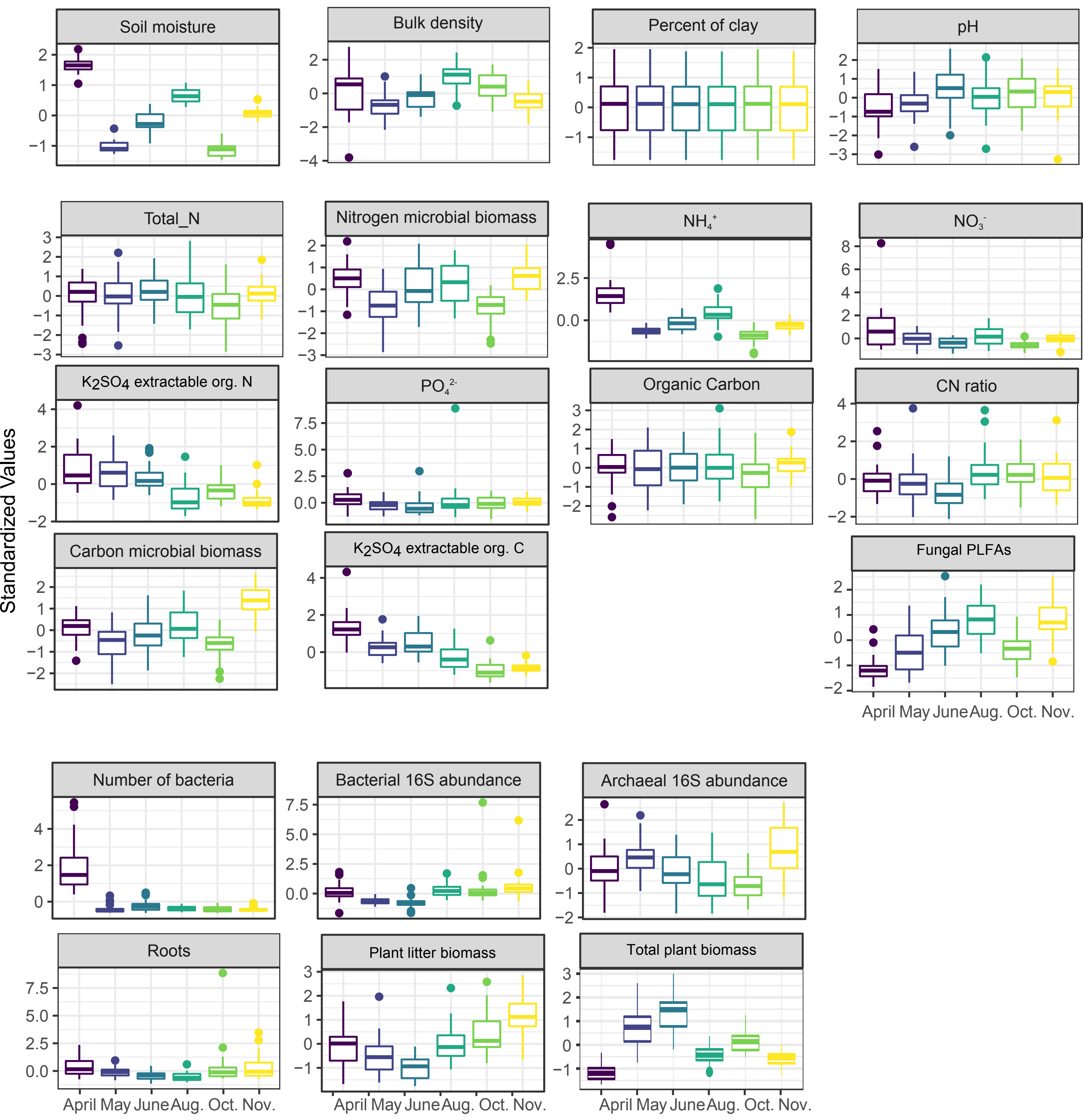


**Figure S2.** Box plots showing the seasonal variation of the environmental parameters from Table S1. Values were normalized (mean=0, SD=1).

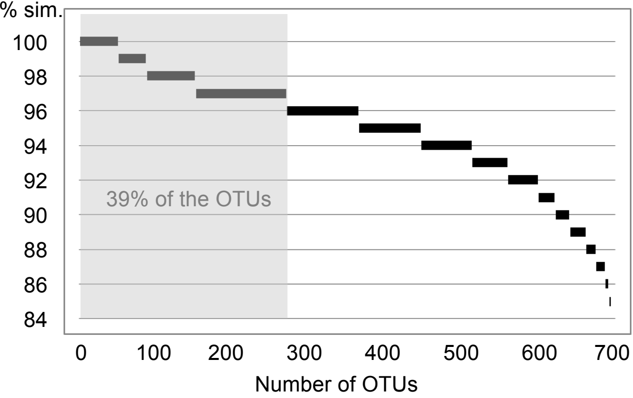


**Figure S3**. Similarities of the OTUs with known sequences. OTUs are classified according to their percentage of similarity to the next kin by BLAST. The horizontal bar length is proportional to the number of OTUs in each rank. Shaded area = OTUs with a similarity ≥ 97%.

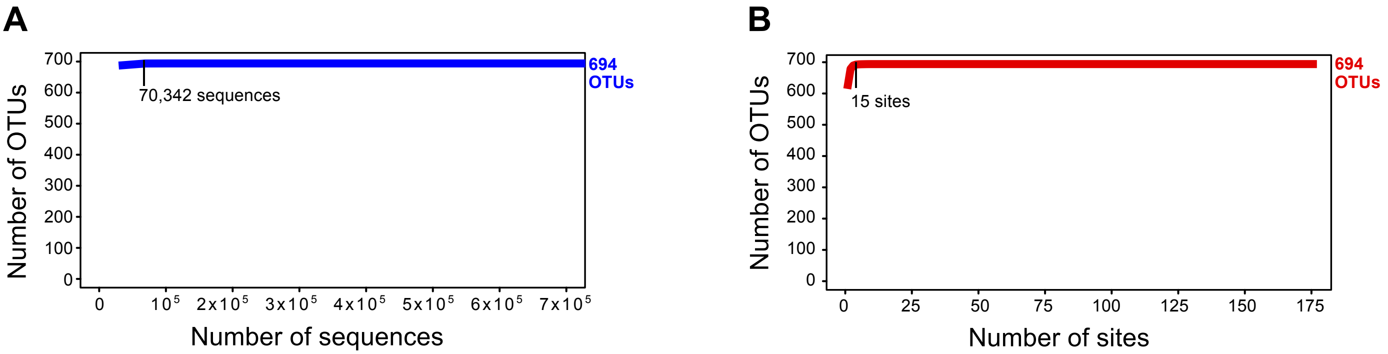


**Figure S4. Description of the diversity. A.** Rarefaction curve describing the observed number of OTUs as a function of the sequencing effort; saturation was reached with c. 70000 sequences**.**

**B.** Species accumulation curve describing the sampling effort; saturation was reached with 15 samples.

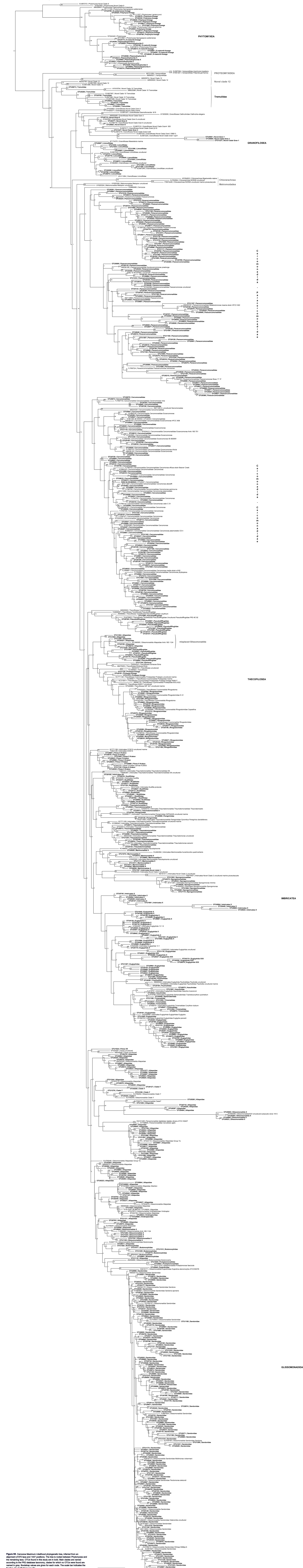


**Figure S5**. Cercozoa Maximum Likelihood phylogenetic tree, inferred from an alignment of 870 taxa and 1447 positions. The tree is rooted between Phytomyxea and the remaining taxa. OTUs found in this study are in bold. Main clades are named according to the PR2 database taxonomy, clades for which no OTUs were found are named in gray. Bootstrap values are given for each node. The scale bar indicates the fraction of substitutions per site.
