## Supplementary material for "Environmental selection and spatiotemporal structure of a major group of soil protists (Rhizaria: Cercozoa) in a temperate grassland": Suppl.TablesS1-S6 Except S3

**Table S1**. Environmental parameters from the study site as in Regan *et al.,* (2014) used in our statistical analyses. Their seasonal variation is shown in Figure S2.

| | Soil_moisture | Bulk_density | Clay_percent | Roots | pH | Total_N | Organic_C | CN_ratio | $NH_4^+$ | $NO_3^-$ | $PO_4^{2-}$ | C_microbial_biomass | N_microbial_biomass | Extractable_Organic_C | Extractable_Organic_N | Fungal_PLFAs | Number_of_bacteria | Bacterial_16S_abundance | Archaeal_16S_abundance | Plant_litter_biomass | Total_plant_biomass |
|---|---|---|---|---|---|---|---|---|---|---|---|---|---|---|---|---|---|---|---|---|---|
| Unit | % | g $cm^{-3}$ | % relative to silt | g/soil core | | µg*g-1 soil dw | µg*g-1 soil dw | | µg*g-1 soil dw | µg*g-1 soil dw | µg*g-1 soil dw | µg*g-1 soil dw | µg*g-1 soil dw | µg*g-1 soil dw | µg*g-1 soil dw | µg g-1 dw | cells g-1 dw | number of copies / g dw | number of copies / g dw | g 20cm2 | g 20cm2 |
| S003 | 63.20 | 0.83 | 16.47 | 2.37 | 6.13 | 0.63 | 6.47 | 10.19 | 13.56 | 24.13 | 88.46 | 1679.34 | 262.90 | 231.08 | 11.99 | 1.54 | 5.64E+10 | 3.96E+11 | 1.14E+09 | 4.98 | 11.28 |
| S004 | 54.50 | 0.79 | 16.06 | 0.87 | 6.49 | 0.56 | 5.51 | 9.84 | 12.36 | 6.59 | 69.07 | 1464.90 | 213.15 | 212.85 | 18.09 | 1.62 | 2.30E+10 | 3.33E+11 | 1.94E+09 | 5.95 | 9.08 |
| S005 | 56.70 | 0.75 | 18.23 | 3.00 | 6.55 | 0.65 | 6.49 | 10.00 | 12.06 | 29.20 | 85.71 | 1787.47 | 271.40 | 200.78 | 8.72 | 1.81 | 3.16E+10 | 4.06E+11 | 3.55E+09 | 2.95 | 8.70 |
| S006 | 59.15 | 0.80 | 17.70 | 3.48 | 6.30 | 0.64 | 6.52 | 10.18 | 14.66 | 22.36 | 83.17 | 1672.75 | 251.27 | 199.55 | 9.04 | 1.83 | 3.13E+10 | 3.24E+11 | 2.48E+09 | 6.61 | 10.89 |
| S007 | 61.24 | 0.77 | 18.10 | 1.22 | 6.52 | 0.68 | 6.89 | 10.20 | 15.13 | 7.09 | 75.90 | 1952.62 | 295.48 | 220.93 | 19.59 | 1.81 | 2.34E+10 | 4.24E+11 | 2.89E+09 | 3.85 | 8.78 |
| S008 | 60.03 | 0.81 | 17.05 | 3.03 | 6.82 | 0.67 | 6.84 | 10.21 | 15.62 | 23.83 | 88.37 | 1874.11 | 293.56 | 220.85 | 10.17 | 1.65 | 2.45E+10 | 3.89E+11 | 2.22E+09 | 5.62 | 9.00 |
| S009 | 61.99 | 0.87 | 16.95 | 1.29 | 6.36 | 0.66 | 6.66 | 10.14 | 16.51 | 22.11 | 72.30 | 1728.85 | 260.16 | 181.07 | 6.95 | 1.98 | 2.39E+10 | 2.12E+11 | 3.79E+09 | 4.72 | 10.38 |
| S010 | 57.33 | 0.74 | 15.70 | 0.90 | 6.57 | 0.68 | 7.16 | 10.59 | 16.87 | 7.70 | 67.68 | 1910.61 | 281.67 | 205.27 | 13.90 | 2.02 | 1.82E+10 | 2.99E+11 | 2.80E+09 | 5.30 | 8.17 |
| S013 | 59.13 | 0.54 | 14.61 | 1.54 | 6.76 | 0.69 | 7.07 | 10.19 | 18.32 | 15.21 | 85.41 | 1790.22 | 311.66 | 253.52 | 11.86 | 1.69 | 4.54E+10 | 4.61E+11 | 5.53E+09 | 2.25 | 4.39 |
| S014 | 54.05 | 0.95 | 15.62 | 1.46 | 6.84 | 0.64 | 6.45 | 10.13 | 13.05 | 24.60 | 93.34 | 1653.95 | 264.78 | 213.66 | 5.67 | 1.60 | 5.42E+10 | 2.58E+11 | 2.19E+09 | 5.26 | 7.12 |
| S015 | 58.75 | 0.98 | 14.67 | 1.62 | 6.56 | 0.63 | 6.32 | 10.11 | 15.91 | 17.61 | 88.30 | 1754.65 | 276.46 | 199.54 | 8.72 | 3.26 | 1.11E+10 | 2.43E+11 | 1.56E+09 | 7.29 | 9.93 |
| S016 | 55.15 | 0.96 | 14.87 | 0.67 | 6.63 | 0.67 | 6.98 | 10.35 | 14.99 | 6.06 | 62.74 | 1755.28 | 257.57 | 212.57 | 16.38 | 2.17 | 3.07E+10 | 3.58E+11 | 3.49E+09 | 3.09 | 6.08 |
| S017 | 56.93 | 0.98 | 14.77 | 1.31 | 6.63 | 0.65 | 6.70 | 10.27 | 19.49 | 24.64 | 67.28 | 1699.59 | 248.41 | 193.71 | 6.28 | 1.85 | 1.69E+10 | 3.62E+11 | 4.16E+07 | 9.20 | 12.17 |
| S018 | 56.71 | 0.99 | 15.09 | 1.70 | 6.55 | 0.65 | 6.49 | 9.94 | 13.93 | 26.82 | 84.85 | 1752.26 | 276.27 | 174.50 | 4.30 | 1.42 | 1.90E+10 | 2.52E+11 | 5.75E+08 | 6.34 | 9.91 |
| S019 | 60.13 | 0.92 | 16.08 | 1.53 | 6.52 | 0.64 | 6.45 | 10.00 | 29.10 | 11.61 | 79.12 | 1820.81 | 280.89 | 355.42 | 32.96 | 2.80 | 2.83E+10 | 3.67E+11 | 2.35E+09 | 3.29 | 8.90 |
| S020 | 58.46 | 1.03 | 15.46 | 1.14 | 6.70 | 0.68 | 6.98 | 10.29 | 13.49 | 26.33 | 73.53 | 1636.69 | 248.00 | 179.65 | 4.27 | 1.59 | 1.09E+10 | 4.07E+11 | 1.76E+09 | 5.55 | 10.19 |
| S021 | 55.72 | 0.97 | 16.20 | 1.08 | 6.51 | 0.67 | 6.69 | 10.03 | 15.48 | 15.15 | 63.55 | 1584.14 | 241.77 | 173.24 | 9.34 | 1.83 | 1.47E+10 | 3.78E+11 | 1.51E+09 | 5.12 | 7.96 |
| S022 | 58.04 | 0.99 | 14.10 | 2.24 | 6.56 | 0.64 | 6.59 | 10.21 | 13.29 | 7.92 | 70.71 | 1605.58 | 246.04 | 191.18 | 13.96 | 1.79 | 3.80E+10 | 6.45E+11 | 2.15E+09 | 4.22 | 6.40 |
| S037 | 57.95 | 0.87 | 12.58 | 2.14 | 7.02 | 0.67 | 6.78 | 10.13 | 17.43 | 7.25 | 101.64 | 1966.03 | 337.32 | 252.74 | 16.32 | 1.99 | 1.29E+10 | 5.30E+11 | 3.37E+09 | 2.46 | 3.47 |
| S038 | 54.59 | 1.00 | 13.54 | 1.21 | 6.80 | 0.62 | 6.23 | 9.97 | 14.17 | 15.57 | 74.56 | 1674.44 | 245.30 | 191.04 | 6.94 | 1.65 | 1.57E+10 | 6.04E+11 | 2.17E+09 | 3.14 | 4.13 |
| S039 | 55.12 | 0.80 | 12.23 | 2.91 | 6.57 | 0.68 | 6.91 | 10.11 | 28.69 | 67.66 | 128.05 | 1970.79 | 296.41 | 225.09 | 6.79 | 1.85 | 2.06E+10 | 2.81E+11 | 2.01E+09 | 2.10 | 3.43 |
| S040 | 57.18 | 1.01 | 12.47 | 2.85 | 6.88 | 0.60 | 5.99 | 9.98 | 18.04 | 12.75 | 128.21 | 1814.46 | 278.28 | 237.06 | 16.62 | 2.36 | 2.93E+10 | 2.45E+11 | 1.76E+09 | 2.91 | 4.01 |
| S041 | 57.45 | 1.04 | 15.05 | 1.98 | 6.75 | 0.66 | 6.55 | 10.00 | 12.12 | 20.79 | 87.29 | 1779.02 | 261.24 | 217.01 | 8.67 | 1.46 | 1.79E+10 | 3.02E+11 | 3.12E+09 | 3.86 | 6.20 |
| S042 | 57.57 | 1.18 | 12.68 | 1.73 | 6.52 | 0.59 | 6.06 | 10.23 | 19.21 | 12.91 | 58.81 | 1456.60 | 215.44 | 187.22 | 9.43 | 1.39 | 1.61E+10 | 1.71E+11 | 1.68E+09 | 4.87 | 6.84 |
| S043 | 57.98 | 1.03 | 13.29 | 1.18 | 6.81 | 0.65 | 6.47 | 9.93 | 15.85 | 6.34 | 73.10 | 1731.53 | 276.13 | 222.94 | 17.99 | 1.85 | 1.34E+10 | 6.77E+09 | 1.67E+09 | 4.39 | 6.15 |
| S044 | 56.62 | 0.96 | 11.83 | 1.74 | 6.70 | 0.67 | 6.60 | 9.92 | 11.67 | 20.54 | 69.75 | 1602.51 | 226.28 | 169.39 | 5.51 | 1.60 | 1.59E+10 | 2.65E+11 | 6.29E+08 | 6.00 | 9.31 |
| S045 | 57.19 | 1.03 | 12.32 | 1.20 | 6.58 | 0.66 | 6.79 | 10.29 | 17.37 | 19.80 | 83.94 | 1895.28 | 308.16 | 187.95 | 6.46 | 2.05 | 2.04E+10 | 3.19E+11 | 3.04E+09 | 0.80 | 2.70 |
| S046 | 58.78 | 0.86 | 13.58 | 2.71 | 6.81 | 0.68 | 7.02 | 10.31 | 17.32 | 6.13 | 94.24 | 1761.92 | 296.48 | 259.85 | 18.11 | 1.75 | 1.19E+10 | 3.10E+11 | 1.56E+09 | 5.35 | 7.36 |
| S049 | 55.00 | 0.99 | 15.13 | 4.07 | 6.55 | 0.57 | 5.74 | 10.06 | 14.41 | 4.74 | 57.87 | 1492.02 | 206.68 | 200.86 | 21.80 | 1.93 | 2.09E+10 | 2.79E+11 | NA | 4.97 | 8.73 |
| S050 | 50.81 | 1.01 | 15.20 | 2.51 | 6.70 | 0.56 | 6.07 | 10.78 | 11.04 | 14.12 | 45.79 | 1343.29 | 191.38 | 141.72 | 3.57 | 1.23 | 1.15E+10 | 2.88E+11 | 1.70E+09 | 5.78 | 7.38 |
| S063 | 26.15 | 0.75 | 16.47 | 2.53 | 6.21 | 0.64 | 6.51 | 10.17 | 7.41 | 7.95 | 59.22 | 1487.35 | 187.18 | 171.99 | 11.23 | 2.27 | 1.02E+10 | 1.42E+11 | 1.52E+09 | 2.14 | 14.20 |
| S064 | 26.83 | 0.85 | 16.06 | 1.06 | 6.68 | 0.61 | 6.08 | 9.93 | 6.61 | 11.72 | 55.04 | 1362.62 | 179.87 | 115.91 | 3.65 | 1.54 | 7.37E+09 | 1.44E+11 | 2.92E+09 | 6.32 | 20.57 |
| S065 | 26.77 | 0.91 | 18.23 | 1.43 | 6.53 | 0.63 | 6.31 | 9.96 | 5.07 | 9.69 | 72.74 | 1362.22 | 181.01 | 160.34 | 12.68 | 1.83 | 2.77E+09 | 1.54E+11 | 2.51E+09 | 2.66 | 7.31 |
| S066 | 34.64 | 0.88 | 17.70 | 0.57 | 6.45 | 0.66 | 6.56 | 9.95 | 5.52 | 12.15 | 81.90 | 1714.88 | 252.90 | 165.02 | 14.35 | 3.24 | 8.04E+09 | 1.76E+11 | 2.76E+09 | 3.36 | 11.44 |
| S067 | 29.43 | 0.84 | 18.10 | 0.66 | 6.65 | 0.64 | 6.47 | 10.12 | 6.74 | 14.44 | 62.78 | 1571.20 | 201.79 | 118.79 | 3.60 | 1.80 | 3.40E+09 | 1.72E+11 | 2.76E+09 | 4.68 | 12.62 |
| S068 | 27.39 | 0.79 | 17.05 | 1.17 | 6.62 | 0.71 | 7.23 | 10.23 | 7.06 | 10.76 | 68.24 | 1554.34 | 222.01 | 152.72 | 9.78 | 3.60 | 5.08E+09 | 1.62E+11 | 3.02E+09 | 4.22 | 13.17 |
| S069 | 29.73 | 0.76 | 16.95 | 2.32 | 6.46 | 0.67 | 6.66 | 9.97 | 5.30 | 7.99 | 76.48 | 1675.64 | 234.34 | 170.64 | 22.88 | 4.10 | 2.08E+09 | 1.77E+11 | 1.74E+09 | 2.42 | 12.91 |
| S070 | 26.13 | 0.78 | 15.70 | 1.97 | 6.62 | 0.69 | 7.14 | 10.41 | 5.57 | 15.49 | 74.25 | 1898.60 | 226.91 | 156.48 | 8.10 | 2.04 | 3.81E+09 | 2.08E+11 | 3.13E+09 | 6.03 | 23.78 |
| S073 | 26.40 | 0.89 | 14.61 | 1.49 | 6.74 | 0.58 | 5.86 | 10.09 | 5.77 | 11.59 | 61.77 | 1676.60 | 187.30 | 123.99 | 6.76 | 1.53 | 2.48E+09 | 1.58E+11 | 2.17E+09 | 2.88 | 7.68 |
| S074 | 27.35 | 0.80 | 15.62 | 1.45 | 6.72 | 0.61 | 6.22 | 10.18 | 6.26 | 8.34 | 60.13 | 1468.12 | 206.80 | 156.43 | 10.50 | 2.18 | 2.82E+09 | 2.04E+11 | 2.42E+09 | 2.26 | 16.38 |
| S075 | 29.64 | 0.92 | 14.67 | 0.74 | 6.50 | 0.64 | 6.47 | 10.09 | 5.37 | 7.80 | 73.10 | 1426.44 | 198.62 | 176.53 | 17.40 | 3.77 | 4.43E+09 | 1.40E+11 | 3.05E+09 | 1.39 | 9.88 |

| | | | | | | | | | | | | | | | | | | | | | |
|---|---|---|---|---|---|---|---|---|---|---|---|---|---|---|---|---|---|---|---|---|---|
| S076 | 26.78 | 0.98 | 14.87 | 0.76 | 6.73 | 0.63 | 6.15 | 9.75 | 6.11 | 14.79 | 60.79 | 1386.20 | 182.36 | 115.04 | 4.18 | 1.38 | 2.37E+09 | 1.49E+11 | 3.19E+09 | 1.88 | 6.99 |
| S077 | 29.24 | 0.87 | 14.77 | 0.88 | 6.58 | 0.61 | 5.96 | 9.83 | 5.68 | 10.31 | 57.39 | 1136.92 | 152.53 | 165.60 | 13.13 | 1.80 | 4.27E+09 | 1.63E+11 | 2.31E+09 | 4.71 | 10.10 |
| S078 | 26.92 | 0.81 | 15.09 | 1.11 | 6.62 | 0.64 | 6.17 | 9.67 | 6.51 | 6.59 | 63.02 | 1640.88 | 242.95 | 168.23 | 13.00 | 2.51 | 5.70E+09 | 1.82E+11 | 2.58E+09 | 4.21 | 13.83 |
| S079 | 25.58 | 0.86 | 16.08 | 1.59 | 6.51 | 0.56 | 5.65 | 10.15 | 6.20 | 2.16 | 46.04 | 1073.71 | 117.31 | 229.82 | 21.23 | 1.63 | 3.33E+09 | 1.05E+11 | 1.14E+09 | 9.67 | 27.61 |
| S080 | 30.36 | 0.87 | 15.46 | 1.03 | 6.70 | 0.70 | 6.95 | 9.99 | 5.94 | 7.64 | 79.27 | 1613.10 | 239.07 | 161.18 | 19.98 | 2.73 | 2.20E+09 | 3.03E+11 | 2.32E+09 | 3.83 | 16.24 |
| S081 | 27.34 | 0.81 | 16.20 | 1.64 | 6.70 | 0.67 | 6.86 | 10.17 | 5.98 | 8.81 | 51.90 | 1695.18 | 239.72 | 150.82 | 10.14 | 2.83 | 2.97E+09 | 2.22E+11 | 3.00E+09 | 3.83 | 14.72 |
| S082 | 27.91 | 0.86 | 14.10 | 1.34 | 6.56 | 0.71 | 7.19 | 10.19 | 5.10 | 13.50 | 59.27 | 1591.73 | 204.17 | 121.09 | 5.44 | 1.88 | 2.90E+09 | 2.16E+11 | 3.23E+09 | 4.00 | 15.32 |
| S097 | 30.08 | 0.74 | 12.58 | 1.60 | 6.83 | 0.67 | 6.58 | 9.90 | 4.30 | 18.72 | 71.90 | 1603.33 | 241.33 | 133.81 | 1.17 | 2.72 | 3.42E+09 | 2.13E+11 | 3.23E+09 | 2.67 | 8.50 |
| S098 | 29.88 | 0.86 | 13.54 | 1.15 | 6.91 | 0.66 | 6.81 | 10.32 | 8.31 | 12.19 | 84.50 | 1416.28 | 212.51 | 181.19 | 9.48 | 2.33 | 1.75E+09 | 2.22E+11 | 3.14E+09 | 1.43 | 12.28 |
| S099 | 26.48 | 0.83 | 12.23 | 1.46 | 6.99 | 0.67 | 7.39 | 11.07 | 7.76 | 9.04 | 92.35 | 1672.98 | 282.92 | 195.87 | 12.53 | 3.13 | 2.87E+09 | 1.33E+11 | 1.48E+09 | 0.96 | 8.26 |
| S100 | 28.30 | 0.95 | 12.47 | 1.60 | 6.87 | 0.66 | 6.93 | 10.49 | 6.31 | 16.15 | 80.04 | 1535.31 | 222.92 | 144.25 | 5.19 | 2.37 | 2.86E+09 | 2.58E+11 | 3.55E+09 | 1.78 | 15.28 |
| S101 | 27.86 | 0.71 | 15.05 | 1.07 | 6.58 | 0.64 | 6.37 | 9.88 | 6.13 | 9.09 | 67.19 | 1192.93 | 160.78 | 157.53 | 17.52 | 1.97 | 2.23E+09 | 2.30E+11 | 2.28E+09 | 3.70 | 12.44 |
| S102 | 26.17 | 0.83 | 12.68 | 2.12 | 6.61 | 0.64 | 6.54 | 10.25 | 7.20 | 5.88 | 71.54 | 1682.17 | 206.23 | 200.22 | 17.82 | 3.43 | 2.95E+09 | 2.18E+11 | 3.76E+09 | 5.65 | 19.13 |
| S103 | 27.21 | 0.86 | 13.29 | 1.05 | 6.79 | 0.72 | 7.41 | 10.25 | 4.26 | 16.91 | 73.18 | 1873.78 | 259.66 | 139.70 | 4.21 | 2.83 | 2.43E+09 | 2.45E+11 | 4.57E+09 | 3.34 | 16.16 |
| S104 | 29.58 | 0.70 | 11.83 | 1.70 | 6.65 | 0.71 | 7.11 | 10.07 | 8.25 | 16.14 | 66.64 | 1461.45 | 236.95 | 148.38 | 7.25 | 2.54 | 3.02E+09 | 2.15E+11 | 4.01E+09 | 4.49 | 13.09 |
| S105 | 30.72 | 0.84 | 12.32 | 1.96 | 6.67 | 0.64 | 6.51 | 10.11 | 6.81 | 11.86 | 68.04 | 1637.90 | 237.39 | 155.25 | 13.05 | 3.15 | 1.83E+09 | 1.96E+11 | 2.72E+09 | 2.33 | 12.97 |
| S106 | 27.56 | 0.77 | 13.58 | 1.75 | 6.75 | 0.67 | 6.71 | 10.05 | 5.46 | 15.66 | 69.24 | 1645.76 | 220.36 | 113.77 | 3.39 | 3.09 | 2.09E+09 | 2.54E+11 | 4.97E+09 | 2.16 | 12.95 |
| S109 | 27.13 | 0.84 | 15.13 | 1.25 | 6.76 | 0.59 | 6.09 | 10.30 | 8.03 | 11.54 | 54.94 | 1693.08 | 205.42 | 127.84 | 7.48 | 2.93 | 2.89E+09 | 2.44E+11 | 1.69E+09 | 6.44 | 15.81 |
| S110 | 26.59 | 1.01 | 15.20 | 0.79 | 6.80 | 0.61 | 6.12 | 10.01 | 5.68 | 4.91 | 47.54 | 1081.53 | 169.12 | 141.45 | 14.09 | 1.80 | 1.83E+09 | 2.30E+11 | 4.55E+09 | 5.90 | 16.55 |
| S123 | 41.89 | 0.92 | 16.47 | 0.72 | 6.72 | 0.69 | 6.88 | 9.95 | 11.79 | 8.78 | 71.47 | 1769.63 | 266.23 | 238.94 | 13.27 | 5.14 | 6.17E+09 | 1.44E+11 | 5.73E+08 | 3.50 | 19.24 |
| S124 | 36.01 | 0.91 | 16.06 | 0.60 | 6.33 | 0.67 | 6.86 | 10.27 | 5.55 | 11.08 | 56.74 | 1626.52 | 226.86 | 145.74 | 10.32 | 3.86 | 5.75E+09 | 9.06E+10 | 1.99E+09 | 3.30 | 18.00 |
| S126 | 36.30 | 0.81 | 17.70 | 0.38 | 6.86 | 0.60 | 5.98 | 10.04 | 7.95 | 7.32 | 47.77 | 1365.01 | 219.47 | 192.81 | 9.19 | 2.41 | 8.27E+09 | 7.73E+10 | 1.84E+09 | 3.60 | 14.20 |
| S127 | 43.53 | 0.83 | 18.10 | 0.74 | 6.89 | 0.70 | 7.32 | 10.42 | 8.27 | 12.92 | 77.64 | 2093.15 | 332.62 | 173.46 | 6.98 | 3.11 | 3.22E+09 | 1.00E+10 | 1.84E+09 | 4.10 | 12.72 |
| S128 | 41.26 | 0.80 | 17.05 | 0.61 | 6.78 | 0.65 | 6.54 | 10.02 | 8.20 | 9.65 | 57.00 | 1639.22 | 259.76 | 155.71 | 7.47 | 3.33 | 3.27E+09 | 3.78E+10 | 2.85E+08 | 3.40 | 17.30 |
| S129 | 40.46 | 0.83 | 16.95 | 0.84 | 7.03 | 0.64 | 6.58 | 10.20 | 10.65 | 12.68 | 63.53 | 1611.04 | 267.44 | 190.08 | 5.49 | 2.52 | 3.07E+09 | 1.36E+11 | 2.10E+09 | 3.20 | 13.60 |
| S130 | 32.91 | 0.93 | 15.70 | 1.26 | 6.57 | 0.64 | 6.57 | 10.22 | 6.62 | 13.26 | 48.76 | 1632.90 | 216.64 | 133.60 | 6.67 | 3.15 | 7.03E+09 | 1.51E+11 | 2.27E+09 | 4.60 | 16.40 |
| S133 | 33.64 | 1.02 | 14.61 | 1.62 | 6.99 | 0.60 | 5.82 | 9.67 | 6.91 | 9.64 | 53.61 | 1576.98 | 237.45 | 153.48 | 5.53 | 3.58 | 6.09E+09 | 1.63E+11 | 2.45E+09 | 0.70 | 11.60 |
| S134 | 35.64 | 0.91 | 15.62 | 0.65 | 7.09 | 0.65 | 6.79 | 10.45 | 9.37 | 11.30 | 81.32 | 1833.31 | 321.50 | 212.16 | 9.52 | 3.20 | 1.05E+10 | 1.46E+11 | 3.64E+09 | 4.50 | 15.20 |
| S135 | 37.85 | 0.92 | 14.67 | 0.72 | 6.84 | 0.64 | 6.29 | 9.81 | 10.05 | 5.04 | 65.23 | 1573.33 | 241.90 | 193.44 | 11.31 | 2.89 | 4.82E+09 | 1.70E+11 | 1.17E+09 | 2.70 | 18.52 |
| S136 | 39.82 | 0.97 | 14.87 | 1.33 | 6.79 | 0.64 | 6.38 | 9.96 | 6.33 | 11.99 | 59.39 | 1602.79 | 238.72 | 126.40 | 6.14 | 2.64 | 5.67E+09 | 2.34E+11 | 1.21E+07 | 2.50 | 13.90 |
| S137 | 35.19 | 0.95 | 14.77 | 1.29 | 6.83 | 0.64 | 6.28 | 9.85 | 6.60 | 8.58 | 61.47 | 1676.15 | 224.98 | 149.63 | 9.57 | 2.94 | 1.17E+10 | 1.50E+11 | 3.33E+09 | 3.30 | 22.80 |
| S138 | 39.97 | 0.86 | 15.09 | 1.99 | 6.82 | 0.64 | 6.16 | 9.64 | 8.97 | 5.01 | 58.65 | 1442.43 | 235.34 | 170.49 | 9.90 | 2.66 | 5.97E+09 | 1.07E+11 | 1.18E+07 | 3.70 | 13.70 |
| S139 | 32.51 | 0.79 | 16.08 | 0.77 | 6.40 | 0.67 | 6.56 | 9.86 | 5.72 | 8.61 | 50.34 | 1431.31 | 181.57 | 125.60 | 7.55 | 1.98 | 8.49E+09 | 1.58E+11 | 3.34E+09 | 2.40 | 19.80 |
| S140 | 29.36 | 0.83 | 15.46 | 0.89 | 6.67 | 0.68 | 6.85 | 10.10 | 6.82 | 5.29 | 53.81 | 1232.05 | 167.23 | 227.34 | 17.71 | 2.14 | 4.37E+09 | 1.32E+11 | 1.82E+09 | 2.40 | 12.13 |
| S141 | 42.96 | 0.91 | 16.20 | 0.24 | 6.96 | 0.67 | 6.84 | 10.17 | 12.13 | 3.70 | 80.82 | 1739.37 | 299.69 | 203.07 | 14.26 | 3.11 | 6.41E+09 | 1.75E+11 | 1.83E+09 | 1.40 | 14.90 |
| S142 | 38.60 | 0.85 | 14.10 | 1.38 | 6.59 | 0.62 | 6.27 | 10.05 | 8.81 | 8.62 | 51.51 | 1518.47 | 214.31 | 129.57 | 7.18 | 3.17 | 4.89E+09 | 1.45E+11 | 1.55E+09 | 3.30 | 13.90 |
| S157 | 36.31 | 0.90 | 12.58 | 1.10 | 7.12 | 0.66 | 6.67 | 10.09 | 5.52 | 11.86 | 132.00 | 1755.40 | 302.07 | 184.10 | 5.66 | 3.54 | 4.08E+09 | 2.09E+11 | 2.10E+09 | 2.60 | 16.10 |
| S158 | 36.15 | 0.80 | 13.54 | 1.10 | 7.23 | 0.68 | 7.01 | 10.35 | 9.11 | 11.63 | 82.23 | 1796.92 | 278.15 | 198.56 | 6.42 | 3.59 | 1.70E+09 | 1.40E+11 | 1.95E+09 | 1.10 | 15.40 |
| S159 | 40.01 | 0.85 | 12.23 | 1.75 | 6.95 | 0.71 | 7.12 | 9.99 | 9.89 | 2.82 | 94.06 | 1831.97 | 328.21 | 223.16 | 17.07 | 3.96 | 4.03E+09 | 1.41E+11 | 1.64E+09 | 1.40 | 17.90 |
| S160 | 36.28 | 0.93 | 12.47 | 1.18 | 7.13 | 0.66 | 6.56 | 9.95 | 7.17 | 10.57 | 71.05 | 1636.66 | 283.51 | 151.42 | 4.41 | 3.55 | 4.51E+09 | 2.87E+11 | 1.40E+09 | 0.60 | 11.90 |
| S161 | 34.51 | 0.77 | 15.05 | 0.96 | 6.73 | 0.65 | 6.33 | 9.74 | 8.77 | 5.91 | 51.21 | 1547.23 | 207.31 | 145.83 | 9.49 | 3.41 | 4.98E+09 | 1.93E+11 | 3.82E+09 | 2.90 | 17.00 |
| S162 | 38.09 | 0.90 | 12.68 | 1.21 | 6.72 | 0.61 | 6.09 | 10.00 | 5.81 | 2.38 | 54.09 | 1397.36 | 211.27 | 158.27 | 12.40 | 2.19 | 2.22E+09 | 2.46E+11 | 2.70E+09 | 2.20 | 16.00 |
| S163 | 33.92 | 0.95 | 13.29 | 0.75 | 6.87 | 0.66 | 6.48 | 9.85 | 5.38 | 9.09 | 55.13 | 1682.44 | 237.88 | 132.48 | 5.94 | 3.69 | 3.87E+09 | 2.04E+11 | 3.99E+09 | 2.70 | 14.98 |
| S164 | 37.93 | 0.83 | 11.83 | 1.19 | 6.82 | 0.66 | 6.53 | 9.86 | 10.55 | 11.12 | 62.54 | 1824.83 | 264.77 | 144.37 | 4.15 | 2.75 | 9.05E+09 | 1.87E+11 | 3.71E+09 | 2.20 | 16.50 |
| S165 | 36.17 | 0.86 | 12.32 | 1.93 | 6.81 | 0.71 | 6.95 | 9.83 | 10.46 | 4.10 | 68.54 | 1806.26 | 304.28 | 202.86 | 18.56 | 3.98 | 2.18E+09 | 1.71E+11 | 2.16E+09 | 1.90 | 19.60 |
| S166 | 37.10 | 0.92 | 13.58 | 1.14 | 6.96 | 0.69 | 6.89 | 9.95 | 7.72 | 11.32 | 62.34 | 1807.64 | 292.30 | 157.75 | 7.69 | 4.40 | 3.57E+09 | 3.96E+11 | 2.85E+09 | 0.70 | 12.30 |
| S169 | 33.97 | 0.91 | 15.13 | 1.06 | 6.61 | 0.60 | 5.78 | 9.66 | 6.63 | 6.09 | 60.71 | 1476.59 | 203.06 | 115.57 | 5.11 | 3.49 | 3.15E+09 | 2.49E+11 | 3.40E+09 | 1.40 | 8.22 |
| S170 | 35.27 | 0.85 | 15.20 | 0.72 | 6.76 | 0.60 | 5.87 | 9.71 | 9.62 | 8.61 | 49.65 | 1370.97 | 204.05 | 123.07 | 2.77 | 2.46 | 2.10E+09 | 1.36E+11 | 1.20E+09 | 0.60 | 16.14 |
| S183 | 44.43 | 1.08 | 16.47 | 0.71 | 6.19 | 0.60 | 6.05 | 10.03 | 10.63 | 8.83 | 44.64 | 1516.20 | 201.70 | 99.58 | 0.32 | 3.21 | 4.59E+09 | 2.56E+11 | 1.68E+09 | 4.01 | 8.95 |

| | | | | | | | | | | | | | | | | | | | |
|---|---|---|---|---|---|---|---|---|---|---|---|---|---|---|---|---|---|---|---|
| S184 | 50.82 | 0.98 | 16.06 | 1.30 | 6.72 | 0.66 | 7.20 | 10.90 | 14.07 | 17.93 | 79.94 | 2000.09 | 313.71 | 143.64 | 2.15 | 3.97 | 2.46E+09 | 3.90E+11 | 2.00E+09 | 3.64 | 9.31 |
| S186 | 46.65 | 0.88 | 17.70 | 1.28 | 6.49 | 0.59 | 5.84 | 9.90 | 6.44 | 10.31 | 45.96 | 1540.71 | 217.82 | 91.85 | 4.95 | 2.92 | 4.70E+09 | 2.63E+11 | 4.08E+09 | 4.73 | 11.14 |
| S187 | 48.51 | 0.84 | 18.10 | 1.34 | 6.80 | 0.67 | 6.73 | 10.07 | 4.69 | 23.30 | 70.64 | 1722.27 | 256.44 | 121.20 | -2.05 | 2.86 | 4.42E+09 | 4.10E+11 | 2.61E+09 | 6.70 | 12.59 |
| S188 | 48.47 | 1.02 | 17.05 | 1.00 | 6.43 | 0.69 | 7.10 | 10.32 | 12.44 | 7.21 | 59.56 | 1467.61 | 208.23 | 174.63 | 9.06 | 2.99 | 3.07E+09 | 3.05E+11 | 2.85E+09 | 5.21 | 12.23 |
| S189 | 51.16 | 0.92 | 16.95 | 1.13 | 6.54 | 0.72 | 7.40 | 10.22 | 9.86 | 16.77 | 67.59 | 1969.24 | 290.92 | 103.57 | -3.06 | 2.62 | 5.76E+09 | 2.79E+11 | 4.10E+09 | 4.66 | 7.00 |
| S190 | 50.20 | 0.94 | 15.70 | 0.38 | 6.83 | 0.74 | 7.81 | 10.51 | 10.44 | 21.98 | 77.59 | 2108.15 | 319.75 | 142.03 | 0.73 | 4.63 | 4.98E+09 | 5.61E+11 | 6.49E+08 | 5.75 | 11.22 |
| S193 | 44.75 | 1.02 | 14.61 | 0.79 | 6.82 | 0.62 | 6.32 | 10.26 | 10.23 | 16.83 | 66.64 | 1709.41 | 238.67 | 116.64 | -0.63 | 3.58 | 2.99E+09 | 3.32E+11 | 2.82E+09 | 6.71 | 13.70 |
| S194 | 44.75 | 1.05 | 15.62 | 0.82 | 6.63 | 0.62 | 6.20 | 10.18 | 11.70 | 7.89 | 79.22 | 1603.50 | 268.49 | 150.13 | 8.24 | 4.10 | 3.99E+09 | 4.56E+11 | 1.48E+09 | 2.74 | 11.77 |
| S195 | 51.01 | 1.04 | 14.67 | 0.60 | 6.66 | 0.70 | 7.21 | 10.37 | 17.25 | 17.44 | 89.34 | 2147.03 | 318.58 | 123.30 | -1.75 | 3.90 | 5.48E+09 | 3.51E+11 | 1.72E+09 | 4.08 | 9.97 |
| S196 | 49.77 | 1.14 | 14.87 | 0.86 | 6.78 | 0.70 | 7.19 | 10.32 | 9.55 | 20.10 | 87.73 | 2041.09 | 308.34 | 135.23 | -1.51 | 4.04 | 5.03E+09 | 4.04E+11 | 3.26E+09 | 5.39 | 11.22 |
| S197 | 48.23 | 1.12 | 14.77 | 0.50 | 6.60 | 0.65 | 6.41 | 10.14 | 12.78 | 8.25 | 65.32 | 1501.40 | 243.20 | 150.24 | 4.15 | 3.56 | 3.10E+09 | 3.84E+11 | 1.28E+09 | 5.62 | 9.47 |
| S198 | 45.78 | 1.00 | 15.09 | 0.76 | 6.63 | 0.66 | 6.55 | 9.93 | 10.30 | 13.67 | 73.89 | 1782.64 | 259.24 | 94.72 | -1.96 | 3.30 | 2.33E+09 | 4.44E+11 | 3.00E+09 | 5.11 | 8.93 |
| S199 | 48.17 | 0.97 | 16.08 | 0.73 | 6.85 | 0.65 | 6.71 | 10.30 | 8.66 | 18.60 | 80.55 | 2003.91 | 288.82 | 106.83 | -4.29 | 4.85 | 2.52E+09 | 5.04E+11 | 2.68E+09 | 7.97 | 14.30 |
| S200 | 48.82 | 1.08 | 15.46 | 0.39 | 6.88 | 0.69 | 7.20 | 10.63 | 15.80 | 7.38 | 99.24 | 1897.84 | 292.64 | 168.49 | 8.01 | 4.83 | 2.59E+09 | 4.16E+11 | 1.61E+09 | 6.71 | 12.79 |
| S201 | 46.32 | 0.86 | 16.20 | 1.52 | 6.64 | 0.64 | 6.68 | 10.38 | 12.41 | 12.22 | 65.74 | 1748.00 | 256.11 | 86.38 | -2.63 | 3.75 | 3.48E+09 | 3.78E+11 | 1.37E+09 | 3.02 | 7.40 |
| S202 | 43.16 | 1.04 | 14.10 | 0.58 | 6.61 | 0.66 | 6.72 | 10.21 | 11.23 | 13.53 | 50.58 | 1670.17 | 217.47 | 116.70 | 0.99 | 4.12 | 2.00E+09 | 3.47E+11 | 1.67E+09 | 3.76 | 10.90 |
| S217 | 49.27 | 0.94 | 12.58 | 0.68 | 7.12 | 0.64 | 6.53 | 10.18 | 11.86 | 19.30 | 89.19 | 1837.16 | 294.63 | 150.45 | -0.94 | 3.31 | 3.67E+09 | 3.49E+11 | 2.03E+08 | 10.55 | 18.00 |
| S218 | 44.07 | 1.08 | 13.54 | 0.92 | 7.14 | 0.67 | 6.83 | 10.16 | 7.76 | 6.14 | 251.29 | 1677.34 | 262.07 | 163.81 | 10.36 | 3.67 | 4.50E+09 | 2.08E+11 | 2.45E+08 | 3.66 | 10.50 |
| S219 | 43.31 | 1.05 | 12.23 | 1.47 | 6.75 | 0.61 | 6.15 | 10.09 | 10.03 | 9.36 | 62.44 | 1645.01 | 236.26 | 83.04 | -1.39 | 2.42 | 5.91E+09 | 3.03E+11 | 1.06E+09 | 2.94 | 9.80 |
| S220 | 44.04 | 1.01 | 12.47 | 1.16 | 6.89 | 0.59 | 6.05 | 10.34 | 9.18 | 12.97 | 66.56 | 1717.80 | 229.02 | 103.67 | -2.02 | 3.61 | 3.21E+09 | 4.22E+11 | 1.33E+09 | 3.59 | 11.92 |
| S221 | 44.19 | 1.07 | 15.05 | 0.39 | 6.61 | 0.61 | 6.30 | 10.32 | 12.11 | 4.26 | 72.96 | 1426.68 | 192.30 | 205.07 | 15.69 | 4.16 | 3.86E+09 | 2.39E+11 | 2.61E+08 | 4.56 | 9.66 |
| S222 | 42.38 | 1.03 | 12.68 | 0.55 | 6.61 | 0.62 | 6.26 | 10.07 | 8.79 | 9.62 | 55.04 | 1568.17 | 219.24 | 85.72 | -0.58 | 2.55 | 2.95E+09 | 2.08E+11 | 5.54E+08 | 4.38 | 10.49 |
| S223 | 44.96 | 0.93 | 13.29 | 1.46 | 6.70 | 0.64 | 6.52 | 10.15 | 9.59 | 12.40 | 63.14 | 1652.38 | 213.58 | 130.61 | 0.41 | 4.13 | 2.40E+09 | 4.15E+11 | 1.38E+09 | 2.98 | 7.68 |
| S224 | 46.29 | 1.03 | 11.83 | 1.21 | 6.73 | 0.64 | 6.52 | 10.15 | 12.02 | 6.67 | 58.74 | 1616.97 | 230.89 | 138.66 | 5.54 | 3.10 | 3.80E+09 | 3.35E+11 | 9.06E+08 | 7.02 | 15.53 |
| S225 | 45.92 | 0.99 | 12.32 | 0.51 | 6.75 | 0.65 | 6.57 | 10.07 | 12.71 | 12.27 | 70.04 | 1730.07 | 259.18 | 111.90 | 0.32 | 3.31 | 6.24E+09 | 2.31E+11 | 5.44E+05 | 4.53 | 9.33 |
| S226 | 46.17 | 1.06 | 13.58 | 0.91 | 6.82 | 0.63 | 6.36 | 10.05 | 10.18 | 13.65 | 58.17 | 1624.59 | 228.49 | 138.03 | -1.95 | 4.42 | 2.96E+09 | 2.90E+11 | 2.50E+09 | 6.51 | 12.57 |
| S229 | 47.06 | 0.99 | 15.13 | 2.14 | 6.81 | 0.62 | 6.42 | 10.42 | 17.31 | 15.13 | 98.81 | 1957.84 | 286.01 | 169.63 | 3.09 | 3.72 | 3.12E+09 | 4.23E+11 | 8.98E+08 | 4.05 | 9.11 |
| S230 | 42.89 | 1.02 | 15.20 | 0.62 | 6.78 | 0.59 | 6.55 | 11.05 | 7.58 | 3.90 | 62.78 | 1386.36 | 184.33 | 115.50 | 5.31 | 2.64 | 5.05E+09 | 6.24E+11 | 3.54E+08 | 2.29 | 7.55 |
| S243 | 29.18 | 1.03 | 16.47 | 0.85 | 6.97 | 0.64 | 6.49 | 10.13 | 7.47 | 8.76 | 86.17 | 1647.80 | 221.67 | 87.68 | 3.90 | 3.49 | 3.91E+09 | 3.23E+11 | 2.17E+09 | 5.90 | 12.35 |
| S244 | 29.27 | 0.95 | 16.06 | 2.12 | 6.61 | 0.68 | 7.29 | 10.67 | 8.48 | 6.07 | 82.29 | 1428.15 | 205.73 | 86.81 | 3.05 | 2.74 | 2.47E+09 | 3.73E+11 | 6.15E+08 | 4.09 | 12.68 |
| S245 | 27.00 | 1.08 | 18.23 | 2.91 | 6.62 | 0.64 | 6.53 | 10.19 | 6.02 | 6.55 | 77.19 | 1608.47 | 211.72 | 88.31 | 3.21 | 2.75 | 2.02E+09 | 4.36E+11 | 2.70E+09 | 6.14 | 14.68 |
| S246 | 26.81 | 0.94 | 17.70 | 1.63 | 6.86 | 0.67 | 6.67 | 10.01 | 5.62 | 6.59 | 82.08 | 1798.20 | 243.73 | 107.38 | 8.24 | 3.12 | 5.41E+09 | 3.18E+11 | 1.64E+09 | 11.18 | 20.99 |
| S247 | 32.91 | 0.89 | 18.10 | 1.17 | 6.72 | 0.66 | 6.64 | 10.14 | 5.61 | 8.05 | 95.14 | 1504.48 | 216.40 | 83.62 | 5.61 | 2.05 | 6.79E+09 | 2.99E+11 | 1.48E+09 | 5.16 | 13.15 |
| S248 | 28.16 | 1.05 | 17.05 | 0.85 | 6.75 | 0.64 | 6.46 | 10.09 | 7.05 | 3.33 | 87.39 | 1608.98 | 210.28 | 97.72 | 7.20 | 3.17 | 6.52E+09 | 2.56E+11 | 3.01E+08 | 9.33 | 20.23 |
| S249 | 24.62 | 0.92 | 16.95 | 0.95 | 6.54 | 0.62 | 6.42 | 10.37 | 4.66 | 6.10 | 71.47 | 1612.19 | 201.81 | 76.84 | 1.34 | 2.58 | 2.71E+09 | 2.84E+11 | 2.25E+09 | 7.15 | 14.90 |
| S250 | 25.91 | 0.98 | 15.70 | 1.37 | 6.57 | 0.60 | 6.17 | 10.22 | 2.00 | 6.46 | 75.14 | 1321.67 | 165.76 | 75.57 | 4.77 | 2.15 | 4.54E+09 | 3.49E+11 | 1.27E+09 | 4.87 | 14.14 |
| S253 | 29.60 | 0.88 | 14.61 | 2.37 | 6.92 | 0.58 | 6.02 | 10.34 | 0.55 | 7.19 | 80.44 | 1522.30 | 208.25 | 72.58 | 6.58 | 2.58 | 2.40E+09 | 2.78E+11 | 1.59E+09 | 8.05 | 16.08 |
| S254 | 27.35 | 0.80 | 15.62 | 1.15 | 7.13 | 0.59 | 5.85 | 9.91 | 5.84 | 7.65 | 62.17 | 1470.21 | 216.02 | 81.90 | 1.12 | 2.68 | 3.24E+09 | 3.66E+11 | 1.10E+09 | 5.92 | 11.49 |
| S255 | 25.90 | 0.96 | 14.67 | 1.04 | 6.93 | 0.61 | 6.31 | 10.35 | 5.08 | 9.96 | 62.75 | 1641.23 | 237.75 | 96.24 | 1.31 | 2.26 | 1.72E+09 | 2.74E+11 | 2.26E+09 | 4.85 | 12.64 |
| S256 | 28.30 | 0.97 | 14.87 | 2.14 | 7.04 | 0.63 | 6.48 | 10.32 | 0.20 | 6.68 | 83.50 | 1520.62 | 226.42 | 89.07 | 12.80 | 1.91 | 1.84E+09 | 5.56E+11 | 2.11E+09 | 6.20 | 12.18 |
| S257 | 24.67 | 0.95 | 14.77 | 1.34 | 6.67 | 0.61 | 6.02 | 9.79 | 4.55 | 6.92 | 68.27 | 1582.19 | 238.92 | 113.20 | 4.01 | 2.02 | 2.90E+09 | 2.52E+11 | 1.61E+09 | 7.79 | 16.26 |
| S258 | 24.82 | 0.99 | 15.09 | 11.15 | 6.88 | 0.58 | 5.82 | 10.04 | 4.36 | 7.12 | 43.34 | 1136.55 | 134.32 | 123.35 | 5.49 | 2.31 | 1.81E+09 | 1.73E+12 | 1.03E+09 | 4.88 | 13.75 |
| S259 | 24.38 | 1.01 | 16.08 | 1.57 | 6.93 | 0.61 | 6.13 | 10.13 | 4.58 | 2.84 | 52.65 | 1516.35 | 194.74 | 92.68 | 7.91 | 2.53 | 4.37E+09 | 2.02E+11 | 5.89E+08 | 7.19 | 13.35 |
| S260 | 23.39 | 0.95 | 15.46 | 0.76 | 6.83 | 0.60 | 5.99 | 10.00 | 3.95 | 2.92 | 41.12 | 1260.46 | 141.72 | 173.97 | 10.41 | 2.15 | 1.62E+09 | 2.15E+11 | 9.14E+08 | 9.06 | 14.88 |
| S261 | 24.01 | 0.93 | 16.20 | 0.77 | 6.69 | 0.68 | 7.19 | 10.61 | 3.47 | 10.45 | 53.66 | 1568.36 | 213.57 | 82.08 | 0.31 | 2.68 | 2.94E+09 | 5.96E+11 | 9.60E+08 | 3.14 | 9.82 |
| S262 | 24.90 | 0.86 | 14.10 | 1.21 | 6.38 | 0.63 | 6.58 | 10.39 | 5.89 | 4.39 | 53.28 | 1471.09 | 194.14 | 75.32 | 3.51 | 2.79 | 2.77E+09 | 3.74E+11 | 1.30E+09 | 4.57 | 12.84 |
| S277 | 28.71 | 0.85 | 12.58 | 2.48 | 7.01 | 0.64 | 6.65 | 10.35 | 4.04 | 5.05 | 86.67 | 1573.03 | 227.84 | 71.75 | 5.82 | 2.15 | 2.46E+09 | 2.83E+11 | 1.67E+09 | 3.08 | 9.48 |
| S278 | 28.48 | 0.94 | 13.54 | 1.80 | 6.94 | 0.66 | 6.86 | 10.39 | 5.57 | 6.02 | 66.00 | 1680.05 | 226.67 | 112.90 | 4.97 | 3.18 | 2.98E+09 | 3.87E+11 | 1.71E+09 | 4.56 | 11.82 |
| S279 | 26.11 | 0.90 | 12.23 | 0.81 | 6.72 | 0.65 | 6.59 | 10.12 | 4.15 | 12.53 | 75.85 | 1626.02 | 231.22 | 77.65 | -0.95 | 2.58 | 3.38E+09 | 3.25E+11 | 2.52E+09 | 4.59 | 13.43 |

| | | | | | | | | | | | | | | | | | | | | |
|---|---|---|---|---|---|---|---|---|---|---|---|---|---|---|---|---|---|---|---|---|
| S280 | 26.36 | 0.82 | 12.47 | 0.90 | 6.48 | 0.58 | 6.02 | 10.37 | 4.69 | 6.49 | 62.02 | 1384.07 | 178.17 | 62.19 | 1.23 | 2.20 | 3.71E+09 | 3.47E+11 | 7.68E+08 | 3.92 | 14.47 |
| S281 | 27.84 | 1.02 | 15.05 | 1.13 | 6.91 | 0.63 | 6.44 | 10.20 | 8.19 | 7.01 | 67.75 | 1529.13 | 212.08 | 113.18 | 6.11 | 2.59 | 4.26E+09 | 3.02E+11 | 1.09E+09 | 4.91 | 11.96 |
| S282 | 24.29 | 1.03 | 12.68 | 1.72 | 6.57 | 0.55 | 5.47 | 10.02 | 2.71 | 7.39 | 50.24 | 1217.72 | 148.02 | 83.71 | 2.37 | 1.56 | 2.76E+09 | 2.95E+11 | 1.78E+09 | 3.77 | 10.47 |
| S283 | 28.00 | 1.01 | 13.29 | 1.20 | 6.90 | 0.66 | 6.95 | 10.50 | 6.55 | 11.53 | 74.55 | 1779.46 | 241.57 | 116.93 | 3.82 | 3.71 | 5.86E+09 | 4.21E+11 | 1.06E+09 | 5.92 | 18.21 |
| S284 | 27.70 | 1.02 | 11.83 | 1.84 | 6.58 | 0.63 | 6.39 | 10.14 | 6.18 | 4.63 | 65.32 | 1578.71 | 193.57 | 109.77 | 4.69 | 2.86 | 4.49E+09 | 2.96E+11 | 7.35E+08 | 2.94 | 11.96 |
| S285 | 27.86 | 0.89 | 12.32 | 0.85 | 6.91 | 0.70 | 7.27 | 10.35 | 5.19 | 11.71 | 90.92 | 1815.99 | 250.00 | 126.60 | 6.95 | 3.20 | 5.18E+09 | 3.65E+11 | 2.12E+08 | 9.83 | 18.97 |
| S286 | 28.17 | 1.02 | 13.58 | 1.62 | 6.75 | 0.62 | 6.28 | 10.15 | 6.64 | 9.09 | 66.80 | 1564.28 | 198.91 | 89.69 | 0.20 | 3.10 | 5.24E+09 | 3.83E+11 | 9.09E+08 | 8.31 | 20.88 |
| S289 | 29.48 | 0.88 | 15.13 | 1.74 | 6.63 | 0.62 | 6.45 | 10.35 | 4.97 | 9.27 | 59.62 | 1483.74 | 190.12 | 74.80 | -0.72 | 2.24 | 2.44E+09 | 3.57E+11 | 3.04E+09 | 5.22 | 12.10 |
| S290 | 23.98 | 0.92 | 15.20 | 3.80 | 6.82 | 0.60 | 6.23 | 10.31 | 5.04 | 2.71 | 78.65 | 1368.17 | 167.63 | 96.53 | 6.05 | 2.43 | 4.89E+09 | 2.82E+11 | 1.88E+09 | 3.45 | 12.65 |
| S303 | 37.03 | 0.97 | 16.47 | 0.91 | 6.67 | 0.60 | 6.27 | 10.39 | 6.41 | 7.97 | 75.24 | 1872.68 | 239.81 | 94.61 | 0.98 | 3.33 | 2.70E+09 | 1.87E+11 | 3.14E+09 | 3.65 | 7.03 |
| S304 | 40.67 | 0.84 | 16.06 | 1.76 | 6.52 | 0.62 | 6.52 | 10.50 | 8.34 | 11.14 | 72.98 | 1848.00 | 238.15 | 92.00 | -0.14 | 2.80 | 2.86E+09 | 3.34E+11 | 2.47E+09 | 7.73 | 13.65 |
| S306 | 38.58 | 0.88 | 17.70 | 1.85 | 6.72 | 0.64 | 6.74 | 10.48 | 7.08 | 11.61 | 76.36 | 2035.63 | 254.66 | 121.55 | 1.88 | 4.15 | 2.43E+09 | 3.78E+11 | 1.87E+09 | 10.35 | 15.36 |
| S307 | 42.81 | 0.73 | 18.10 | 1.12 | 6.52 | 0.64 | 6.44 | 10.01 | 6.92 | 14.02 | 68.98 | 1677.64 | 242.76 | 133.81 | 4.66 | 2.13 | 4.05E+09 | 4.05E+11 | 4.69E+09 | 6.91 | 9.94 |
| S308 | 38.76 | 0.82 | 17.05 | 1.26 | 6.69 | 0.68 | 6.99 | 10.31 | 5.34 | 10.30 | 80.14 | 1939.97 | 243.74 | 93.41 | 3.88 | 3.27 | 2.72E+09 | 4.94E+11 | 2.92E+09 | 6.64 | 11.75 |
| S309 | 39.29 | 0.89 | 16.95 | 1.52 | 6.50 | 0.63 | 6.35 | 10.03 | 10.56 | 13.56 | 69.73 | 2104.29 | 255.34 | 93.56 | -1.99 | 5.12 | 4.91E+09 | 3.90E+11 | 5.06E+09 | 8.42 | 14.19 |
| S310 | 37.22 | 0.83 | 15.70 | 0.68 | 6.63 | 0.66 | 6.71 | 10.20 | 9.61 | 8.93 | 54.31 | 2042.22 | 233.05 | 107.03 | 4.42 | 2.82 | 2.67E+09 | 4.16E+11 | 2.15E+09 | 6.66 | 10.87 |
| S313 | 45.20 | 0.85 | 14.61 | 0.74 | 6.64 | 0.65 | 6.77 | 10.38 | 7.43 | 14.96 | 91.14 | 2189.72 | 326.35 | 109.18 | -0.82 | 3.43 | 3.67E+09 | 3.07E+11 | 5.60E+09 | 7.34 | 13.59 |
| S314 | 40.17 | 0.95 | 15.62 | 2.82 | 7.03 | 0.66 | 6.58 | 9.94 | 6.93 | 7.79 | 77.06 | 1999.65 | 280.05 | 100.29 | 6.53 | 5.13 | 3.34E+09 | 3.40E+11 | 2.00E+09 | 6.44 | 10.69 |
| S315 | 40.96 | 0.87 | 14.67 | 2.59 | 6.85 | 0.66 | 6.73 | 10.24 | 7.93 | 12.86 | 70.45 | 2280.13 | 331.84 | 99.52 | -1.01 | 4.07 | 5.97E+09 | 2.00E+11 | 4.04E+09 | 4.30 | 8.91 |
| S316 | 37.69 | 0.92 | 14.87 | 1.85 | 6.82 | 0.67 | 6.90 | 10.36 | 5.28 | 14.89 | 75.04 | 2099.74 | 278.23 | 104.29 | -0.77 | 2.92 | 3.24E+09 | 4.71E+11 | 2.32E+09 | 3.29 | 5.83 |
| S317 | 40.20 | 0.83 | 14.77 | 3.76 | 6.56 | 0.64 | 6.31 | 9.90 | 7.99 | 10.45 | 55.24 | 1797.80 | 219.86 | 90.96 | 1.66 | 3.73 | 3.00E+09 | 2.73E+11 | 3.56E+09 | 7.62 | 12.18 |
| S318 | 40.39 | 0.92 | 15.09 | 1.88 | 6.08 | 0.63 | 6.14 | 9.82 | 5.03 | 13.88 | 67.28 | 1937.34 | 252.22 | 105.98 | -2.08 | 3.97 | 3.97E+09 | 3.36E+11 | 8.44E+08 | 9.03 | 16.26 |
| S319 | 41.18 | 0.77 | 16.08 | 1.21 | 6.96 | 0.66 | 6.66 | 10.14 | 9.03 | 11.07 | 68.05 | 2193.35 | 284.19 | 100.60 | -1.01 | 3.88 | 3.16E+09 | 3.26E+11 | 4.57E+09 | 8.97 | 13.50 |
| S320 | 40.32 | 0.82 | 15.46 | 5.28 | 6.80 | 0.66 | 6.73 | 10.17 | 8.54 | 3.36 | 66.48 | 2015.93 | 242.09 | 119.61 | 12.93 | 3.96 | 3.11E+09 | 3.59E+11 | 2.81E+09 | 7.93 | 14.57 |
| S321 | 39.47 | 0.86 | 16.20 | 1.07 | 6.86 | 0.65 | 6.66 | 10.26 | 8.36 | 13.57 | 83.66 | 2209.96 | 314.32 | 107.62 | -0.39 | 3.51 | 6.57E+09 | 3.67E+11 | 4.79E+09 | 7.30 | 12.75 |
| S322 | 39.47 | 0.85 | 14.10 | 0.92 | 6.76 | 0.67 | 6.67 | 9.98 | 8.85 | 12.16 | 61.64 | 2075.82 | 265.82 | 103.38 | 1.86 | 3.46 | 2.27E+09 | 3.53E+11 | 3.09E+09 | 6.66 | 12.17 |
| S337 | 38.98 | 0.90 | 12.58 | 2.01 | 7.00 | 0.66 | 6.56 | 9.91 | 6.85 | 12.08 | 83.09 | 2344.36 | 325.24 | 108.55 | -1.13 | 3.28 | 2.79E+09 | 3.93E+11 | 4.34E+09 | 9.75 | 17.09 |
| S338 | 41.02 | 0.80 | 13.54 | 1.03 | 6.81 | 0.64 | 6.61 | 10.39 | 4.94 | 11.12 | 83.44 | 2154.70 | 295.86 | 107.68 | 0.39 | 4.03 | 3.09E+09 | 4.88E+11 | 3.12E+09 | 8.75 | 15.36 |
| S339 | 38.54 | 0.88 | 12.23 | 1.00 | 6.75 | 0.63 | 6.87 | 10.92 | 6.97 | 7.06 | 83.87 | 2130.89 | 268.74 | 88.13 | 2.39 | 3.83 | 5.06E+09 | 4.27E+11 | 3.62E+09 | 10.64 | 17.93 |
| S340 | 39.00 | 0.99 | 12.47 | 0.87 | 6.79 | 0.68 | 7.01 | 10.23 | 8.06 | 9.20 | 71.73 | 2161.65 | 287.39 | 88.47 | -0.21 | 3.05 | 2.85E+09 | 4.57E+11 | 2.16E+09 | 11.32 | 16.42 |
| S341 | 45.06 | 0.81 | 15.05 | 3.69 | 6.78 | 0.66 | 6.67 | 10.09 | 7.34 | 11.62 | 93.06 | 2110.65 | 281.62 | 97.23 | 0.36 | 4.19 | 2.62E+09 | 2.95E+11 | 5.29E+09 | 7.21 | 11.75 |
| S342 | 39.68 | 0.89 | 12.68 | 2.32 | 6.48 | 0.65 | 6.48 | 9.98 | 7.76 | 9.65 | 64.34 | 1828.41 | 236.13 | 78.66 | 1.19 | 4.27 | 4.19E+09 | 6.36E+11 | 1.87E+09 | 8.82 | 14.30 |
| S343 | 42.09 | 0.94 | 13.29 | 1.00 | 6.81 | 0.61 | 6.17 | 10.13 | 6.47 | 10.29 | 69.66 | 1933.12 | 253.28 | 108.84 | -0.88 | 2.88 | 3.17E+09 | 4.33E+11 | 5.28E+09 | 8.04 | 13.07 |
| S344 | 39.68 | 0.91 | 11.83 | 4.52 | 6.92 | 0.67 | 6.79 | 10.13 | 8.46 | 7.76 | 63.08 | 1861.83 | 240.02 | 95.63 | 0.04 | 3.36 | 3.21E+09 | 1.45E+12 | 1.61E+09 | 11.84 | 17.36 |
| S345 | 41.61 | 0.86 | 12.32 | 1.06 | 6.58 | 0.64 | 6.53 | 10.17 | 9.66 | 10.67 | 80.98 | 1878.53 | 269.91 | 86.00 | 0.14 | 3.96 | 3.24E+09 | 4.70E+11 | 4.32E+09 | 5.43 | 11.27 |
| S346 | 40.55 | 0.79 | 13.58 | 1.43 | 6.84 | 0.71 | 7.31 | 10.30 | 8.29 | 15.51 | 74.41 | 2013.37 | 277.48 | 98.62 | -0.78 | 3.40 | 3.38E+09 | 3.94E+11 | 2.46E+09 | 7.03 | 11.16 |
| S349 | 41.60 | 0.85 | 15.13 | 2.63 | 7.01 | 0.65 | 6.46 | 9.94 | 7.84 | 13.01 | 70.60 | 2264.85 | 320.86 | 107.57 | -0.17 | 2.42 | 3.38E+09 | 4.52E+11 | 4.22E+09 | 9.25 | 15.28 |
| S350 | 40.58 | 0.88 | 15.20 | 1.32 | 6.84 | 0.64 | 6.70 | 10.44 | 6.37 | 10.55 | 73.31 | 1999.10 | 273.05 | 100.06 | 1.00 | 4.33 | 3.62E+09 | 4.26E+11 | 2.30E+09 | 6.60 | 14.32 |
| average | 39.36 | 0.91 | 14.87 | 1.49 | 6.72 | 0.65 | 6.55 | 10.16 | 8.99 | 11.34 | 71.96 | 1693.97 | 241.91 | 142.91 | 6.49 | 2.88 | 7.26E+09 | 3.11E+11 | 2.27E+09 | 4.89 | 12.51 |
| minimum | 23.39 | 0.54 | 11.83 | 0.24 | 6.08 | 0.55 | 5.47 | 9.64 | 0.20 | 2.16 | 41.12 | 1073.71 | 117.31 | 62.19 | -4.29 | 1.23 | 1.62E+09 | 6.77E+09 | 5.44E+05 | 0.60 | 2.70 |
| maximum | 63.20 | 1.18 | 18.23 | 11.15 | 7.23 | 0.74 | 7.81 | 11.07 | 29.10 | 67.66 | 251.29 | 2344.36 | 337.32 | 355.42 | 32.96 | 5.14 | 5.64E+10 | 1.73E+12 | 5.60E+09 | 11.84 | 27.61 |
| st. deviation | 10.93 | 0.10 | 1.73 | 1.09 | 0.20 | 0.03 | 0.41 | 0.24 | 4.41 | 6.82 | 20.19 | 247.70 | 43.58 | 49.19 | 6.30 | 0.89 | 9.01E+09 | 1.85E+11 | 1.23E+09 | 2.44 | 4.10 |

*Fiore-Donno et al. 2019* . **Community structure of soil protists (Cercozoa)**

**Table S2**. Combinations of barcodes used in this study, with the corresponding samples.

| Forward | Reverse | Sample | Forward | Reverse | Sample | Forward | Reverse | Sample |
|---|---|---|---|---|---|---|---|---|
| CTACATCT | ACACGACG | S003 | ACACAGCA | GAGTGACG | S075 | GAGCTAGT | AGATCAGC | S040 |
| CAGCATGA | ACACGACG | S004 | CTACATCT | GCATGCGT | S076 | GAGACATA | GAGATCAT | S099 |
| CAGATGTC | ACACGACG | S063 | CAGCATGA | GCATGCGT | S135 | CTGCTGCA | GCGTCGAC | S100 |
| ATGACTCT | ACACGACG | S064 | CAGATGTC | GCATGCGT | S136 | CTACTAGC | TAGTCGCT | S159 |
| ATAGCAGT | ACACGACG | S123 | ATGACTCT | GCATGCGT | S195 | CTACTAGC | AGACTGAC | S160 |
| ATAGAGTC | ACACGACG | S124 | ATAGCAGT | GCATGCGT | S196 | GCAGATGC | GAGTGACG | S219 |
| ACGATACA | ACACGACG | S183 | ATAGAGTC | GCATGCGT | S255 | GCACTATA | TAGCGCAG | S220 |
| ACACAGCA | ACACGACG | S184 | ACGATACA | GCATGCGT | S256 | GCACTATA | ACGAGCGC | S279 |
| CTACATCT | ACGAGCGC | S243 | ACACAGCA | GCATGCGT | S315 | GAGCTAGT | AGCATACT | S280 |
| CAGCATGA | ACGAGCGC | S244 | CTACATCT | GCGTCGAC | S316 | GAGACATA | GAGTGACG | S339 |
| CAGATGTC | ACGAGCGC | S303 | CAGCATGA | GCGTCGAC | S017 | CTGCTGCA | TACATGCG | S340 |
| ATGACTCT | ACGAGCGC | S304 | CAGATGTC | GCGTCGAC | S018 | CTGCTGCA | ACACGACG | S041 |
| ATAGCAGT | ACGAGCGC | S005 | ATGACTCT | GCGTCGAC | S077 | CTACTAGC | AGATCAGC | S042 |
| ATAGAGTC | ACGAGCGC | S006 | ATAGCAGT | GCGTCGAC | S078 | GCAGATGC | GCATGCGT | S101 |
| ACGATACA | ACGAGCGC | S065 | ATAGAGTC | GCGTCGAC | S137 | GCACTATA | TAGTCGCT | S102 |
| ACACAGCA | ACGAGCGC | S066 | ACGATACA | GCGTCGAC | S138 | GCACTATA | AGACTGAC | S161 |
| CAGCATGA | AGACTGAC | S126 | ACACAGCA | GCGTCGAC | S197 | GAGCTAGT | AGCTGCAC | S162 |
| ATGACTCT | AGACTGAC | S186 | CTACATCT | TACATGCG | S198 | GAGACATA | GCATGCGT | S221 |
| ATAGCAGT | AGACTGAC | S245 | CAGCATGA | TACATGCG | S257 | CTGCTGCA | TAGCGCAG | S222 |
| ATAGAGTC | AGACTGAC | S246 | CAGATGTC | TACATGCG | S258 | CTGCTGCA | ACGAGCGC | S281 |
| ACACAGCA | AGACTGAC | S306 | ATGACTCT | TACATGCG | S317 | GCAGATGC | GCGTCGAC | S282 |
| CTACATCT | AGATCAGC | S007 | ATAGCAGT | TACATGCG | S318 | GCACTATA | TCGCTAGT | S341 |
| CAGCATGA | AGATCAGC | S008 | ATAGAGTC | TACATGCG | S019 | GCACTATA | AGATCAGC | S342 |
| CAGATGTC | AGATCAGC | S067 | ACGATACA | TACATGCG | S020 | GAGCTAGT | GAGATCAT | S043 |
| ATGACTCT | AGATCAGC | S068 | ACACAGCA | TACATGCG | S079 | GAGACATA | GCGTCGAC | S044 |
| ATAGCAGT | AGATCAGC | S127 | CTACATCT | TAGCGCAG | S080 | CTGCTGCA | TAGTCGCT | S103 |
| ATAGAGTC | AGATCAGC | S128 | CAGCATGA | TAGCGCAG | S139 | CTGCTGCA | AGACTGAC | S104 |
| ACGATACA | AGATCAGC | S187 | CAGATGTC | TAGCGCAG | S140 | CTACTAGC | AGCTGCAC | S163 |
| ACACAGCA | AGATCAGC | S188 | ATGACTCT | TAGCGCAG | S199 | GCAGATGC | TACATGCG | S164 |
| CTACATCT | AGCATACT | S247 | ATAGCAGT | TAGCGCAG | S200 | GCAGATGC | ACACGACG | S223 |
| CAGCATGA | AGCATACT | S248 | ATAGAGTC | TAGCGCAG | S259 | GCACTATA | AGCATACT | S224 |
| CAGATGTC | AGCATACT | S307 | ACGATACA | TAGCGCAG | S260 | GAGCTAGT | GAGTGACG | S283 |
| ATGACTCT | AGCATACT | S308 | CTACATCT | TAGTCGCT | S319 | GAGACATA | TACATGCG | S284 |
| ATAGCAGT | AGCATACT | S009 | CAGCATGA | TAGTCGCT | S320 | GAGACATA | ACACGACG | S343 |
| ATAGAGTC | AGCATACT | S010 | CAGATGTC | TAGTCGCT | S021 | CTGCTGCA | AGATCAGC | S344 |
| ACGATACA | AGCATACT | S069 | ATGACTCT | TAGTCGCT | S022 | CTACTAGC | GAGATCAT | S045 |
| ACACAGCA | AGCATACT | S070 | ATAGCAGT | TAGTCGCT | S081 | GCAGATGC | TAGCGCAG | S046 |
| CTACATCT | AGCTGCAC | S129 | ATAGAGTC | TAGTCGCT | S082 | GCAGATGC | ACGAGCGC | S105 |
| CAGCATGA | AGCTGCAC | S130 | ACGATACA | TAGTCGCT | S141 | GCACTATA | AGCTGCAC | S106 |
| CAGATGTC | AGCTGCAC | S189 | ACACAGCA | TAGTCGCT | S142 | GAGCTAGT | GCATGCGT | S165 |
| ATGACTCT | AGCTGCAC | S190 | GCAGATGC | AGCATACT | S201 | GAGACATA | TAGCGCAG | S166 |
| ATAGCAGT | AGCTGCAC | S249 | GCACTATA | GCATGCGT | S202 | GAGACATA | ACGAGCGC | S225 |
| ATAGAGTC | AGCTGCAC | S250 | GAGCTAGT | TAGCGCAG | S261 | CTGCTGCA | AGCATACT | S226 |
| ACGATACA | AGCTGCAC | S309 | GAGCTAGT | ACGAGCGC | S262 | CTACTAGC | GAGTGACG | S285 |
| CTACATCT | GAGATCAT | S310 | GAGACATA | AGCATACT | S321 | GCAGATGC | TAGTCGCT | S286 |
| CAGCATGA | GAGATCAT | S013 | CTGCTGCA | GAGTGACG | S322 | GCAGATGC | AGACTGAC | S345 |
| CAGATGTC | GAGATCAT | S014 | CTACTAGC | TACATGCG | S037 | GCACTATA | GAGATCAT | S346 |
| ATGACTCT | GAGATCAT | S073 | CTACTAGC | ACACGACG | S038 | GAGCTAGT | GCGTCGAC | S049 |
| ATAGCAGT | GAGATCAT | S074 | GCAGATGC | AGCTGCAC | S097 | GAGACATA | TAGTCGCT | S050 |
| ATAGAGTC | GAGATCAT | S133 | GCACTATA | GCGTCGAC | S098 | GAGACATA | AGACTGAC | S109 |
| ACGATACA | GAGATCAT | S134 | GAGCTAGT | TAGTCGCT | S157 | CTGCTGCA | AGCTGCAC | S110 |
| ACACAGCA | GAGATCAT | S193 | GAGCTAGT | AGACTGAC | S158 | CTACTAGC | GCATGCGT | S169 |
| CTACATCT | GAGTGACG | S194 | GAGACATA | AGCTGCAC | S217 | GCAGATGC | TCGCTAGT | S170 |
| CAGCATGA | GAGTGACG | S253 | CTGCTGCA | GCATGCGT | S218 | GCAGATGC | AGATCAGC | S229 |
| CAGATGTC | GAGTGACG | S254 | CTACTAGC | TAGCGCAG | S277 | CTACATCT | GAGTGACG | S230 |
| ATGACTCT | GAGTGACG | S313 | CTACTAGC | ACGAGCGC | S278 | GAGCTAGT | TACATGCG | S289 |
| ATAGCAGT | GAGTGACG | S314 | GCAGATGC | GAGATCAT | S337 | GAGCTAGT | ACACGACG | S290 |
| ATAGAGTC | GAGTGACG | S015 | GCACTATA | TACATGCG | S338 | GAGACATA | AGATCAGC | S349 |
| ACGATACA | GAGTGACG | S016 | GCACTATA | ACACGACG | S039 | CTGCTGCA | GAGATCAT | S350 |
| CTGCTGCA | TCGCTAGT | Mock Cerco | | | | | | |

| SUB-PHYLUM | CLASS | ORDER | FAMILY | GENUS | NUTRITION | MORPHOLOGY | LOCOMOTION | REFERENCES |
|---|---|---|---|---|---|---|---|---|
| Endomyxa | | | | | N/A | N/A | N/A | |
| | Phytomyxea[1] | | | | N/A | N/A | N/A | |
| | | Plasmodiophorida | | | N/A | N/A | N/A | |
| | | | Incertae sedis | *Woronina* | parasite (not plant) | flagellate (intracellular) | non motile endoparasite | Neuhauser et al. 2014 |
| | | | *Polymyxa* -lineage | *Ligniera, Polymyxa* | plant parasite | flagellate (intracellular) | non motile endoparasite | Neuhauser et al. 2014 |
| | | | *Spongospora* -lineage | *Spongospora* | plant parasite | flagellate (intracellular) | non motile endoparasite | Neuhauser et al. 2014 |
| Filosa | | | | | N/A | N/A | N/A | |
| | Novel clade 10 | | | | N/A | N/A | N/A | |
| | | Tremulida | | | N/A | N/A | N/A | |
| | | | Tremulidae | *Tremula* | N/A | naked flagellate | creeping/gliding on substrate | Howe et al. 2011 |
| | Novel clade 12 | | | | N/A | N/A | N/A | Bass et al. 2009 |
| | Incertae sedis | | | *Kraken* | bacterivore | naked amoeba | creeping/gliding on substrate | Dumack et al. 2016 |
| | Granofilosea | | | | N/A | N/A | N/A | |
| | | | Novel_Clade_Gran-1-2-3-4-5-6 | | N/A | N/A | N/A | Bass et al. 2009 |
| | | | Massisteriidae | *Massisteria* | bacterivore | naked amoeboflagellate | creeping/gliding on substrate[1] | Mylnikov et al. 2015 |
| | | Cryptofilida | Mesofilidae | *Mesofila* | bacterivore | naked amoeboflagellate | creeping/gliding on substrate[1] | Bass et al. 2009 |
| | | Limnofilida | Limnofilidae | *Limnofila* | bacterivore | naked amoeboflagellate | creeping/gliding on substrate | Howe et al. 2011 |
| | Imbricatea | | | | N/A | N/A | N/A | |
| | | Novel_Clade_3 | | | N/A | N/A | N/A | |
| | | Euglyphida | | | N/A | testate amoeboflagellate/ amoeba (silica)[2] | N/A | Meisterfeld 2000 |
| | | | Assulinidae | *Assulina* | omnivore | testate amoeboflagellate/ amoeba (silica) | creeping/gliding on substrate | Meisterfeld 2000 |
| | | | Euglyphidae | | omnivore[3] | testate amoeboflagellate/ amoeba (silica) | creeping/gliding on substrate | Meisterfeld 2000 |
| | | | | *Euglypha* | omnivore | testate amoeboflagellate/ amoeba (silica) | creeping/gliding on substrate | Meisterfeld 2000 |
| | | | Paulinellidae | *Paulinella* | N/A | testate amoeboflagellate/ amoeba (silica) | creeping/gliding on substrate | Meisterfeld 2000 |
| | | | Sphenoderiidae | *Trachelocorythion, Sphenoderia* | omnivore | testate amoeboflagellate/ amoeba (silica) | creeping/gliding on substrate | Meisterfeld 2000 |
| | | | Trinematidae | *Corythion* | omnivore | testate amoeboflagellate/ amoeba (silica) | creeping/gliding on substrate | Meisterfeld 2000 |
| | | Incertae sedis Imbricatea | | *Discomonas* | N/A | naked flagellate | creeping/gliding on substrate | Chantangsi et al. 2010 |
| | | | Nudifilidae | | N/A | naked amoeboflagellate | N/A | Howe et al. 2011 |
| | | | | *Nudifila* | bacterivore | naked amoeboflagellate | creeping/gliding on substrate | Howe et al. 2011 |
| | | Marimonadida | | | N/A | naked amoeboflagellate | N/A (gliding or swimming) | Howe et al. 2011 |
| | | Spongomonadida | | | N/A | N/A | N/A | |
| | | | Spongomonadidae | | bacterivore | N/A | N/A | Howe et al. 2011; Hibberd 1976; Strüder-Kypke et al. 1998 |
| | | | | *Spongomonas* | bacterivore | naked flagellate | creeping/gliding on substrate[4] | Howe et al. 2011; Hibberd 1976; Strüder-Kypke et al. 1998 |
| | | Thaumatomonadida | | | N/A | N/A | N/A | |
| | | | Peregriniidae | *Peregrinia* | bacterivore | naked amoeboflagellate | creeping/gliding on substrate | Howe et al. 2011 |
| | | | Thaumatomonadidae | | N/A | N/A | N/A | |

| | | | | | | | |
|---|---|---|---|---|---|---|---|
| | | | *Penardeugenia* | omnivore | testate amoeboflagellate/ amoeba (silica) | creeping/gliding on substrate | Dumack et al. 2018 |
| | | | *Thaumatomonas, Thaumatomastix* | bacterivore | testate amoeboflagellate/ amoeba (silica) | creeping/gliding on substrate | Howe et al. 2011 |
| | | | *Reckertia* | bacterivore | testate amoeboflagellate/ amoeba (silica) | creeping/gliding on substrate | Howe et al. 2011 |
| | | | *Esquamula* | bacterivore | naked amoeboflagellate | creeping/gliding on substrate | Shiratori et al. 2012 |
| Sarcomonadea | | | | N/A | N/A | N/A | |
| | Cercomonadida | | | N/A | N/A | N/A | Bass et al. 2009[5] |
| | | Cercomonadidae | | N/A | N/A | N/A | Bass et al. 2009[5] |
| | | | *Cavernomonas* | bacterivore | naked flagellate | creeping/gliding on substrate | Bass et al. 2009 |
| | | | *Cercomonas, Eocercomonas, Cholamonas* | omnivore | naked amoeboflagellate | creeping/gliding on substrate | Ekelund 1998; Flavin et al. 2000; Geisen et al. 2016 |
| | | Paracercomonadidae | | bacterivore | naked amoeboflagellate | N/A | |
| | | | *Metabolomonas* | bacterivore | naked amoeboflagellate | creeping/gliding on substrate | Brabender et al. 2012 |
| | | | *Nucleocercomonas* | bacterivore | naked amoeboflagellate | freely swimming | Brabender et al. 2012 |
| | | | *Paracercomonas* | bacterivore | naked amoeboflagellate | creeping/gliding on substrate | Bass et al. 2009 |
| | Glissomonadida | | | N/A | N/A | N/A | |
| | | Allapsidae | Group-Te, *Allantion, Allapsa, Teretomonas* | bacterivore | naked flagellate | creeping/gliding on substrate | Howe et al. 2009 |
| | | Viridiraptoridae | *Viridiraptor* | eukaryvore | naked amoeboflagellate | creeping/gliding on substrate[1] | Hess et al. 2013 |
| | | Bodomorphidae | *Bodomorpha* | bacterivore | naked flagellate | creeping/gliding on substrate | Howe et al. 2009 |
| | | Clade-T | | N/A | N/A | N/A | Howe et al. 2009 |
| | | Clade-Y | | N/A | N/A | N/A | Howe et al. 2009 |
| | | Dujardinidae | *Dujardina* | bacterivore | naked flagellate | creeping/gliding on substrate | Howe et al. 2009 |
| | | Proleptomonadidae | *Proleptomonas* | bacterivore[6] | naked flagellate | freely swimming | Vickerman et al. 2002 |
| | | Sandonidae | *Clade-N/F-A, Flectomonas, Mollimonas, Neoheteromita, Sandona* | bacterivore | naked flagellate | creeping/gliding on substrate | Howe et al. 2009 |
| | Pansomonadida | | | N/A | naked amoeboflagellate | creeping/gliding on substrate[1] | Howe et al. 2011 |
| | | Agitatidae | | N/A | naked amoeboflagellate | creeping/gliding on substrate[1] | Howe et al. 2011 |
| | | | *Aurigamonas* | omnivore | naked amoeboflagellate | creeping/gliding on substrate[1] | Vieckerman et al. 2005 |
| | | | *Agitata* | bacterivore | naked amoeboflagellate | creeping/gliding on substrate[1] | Howe et al. 2011 |
| Thecofilosea | | | | N/A | N/A | N/A | |
| | Cryomonadida | | | N/A | testate amoeboflagellate/ amoeba/flagellate (organic/agglutinated) | N/A | Cavalier-Smith 1993 |
| | | *Cryomonas* -lineage | *Cryothecomonas* | eukaryvore | testate amoeboflagellate/ amoeba/flagellate (organic/agglutinated) | N/A | Thomsen et al. 1991 |
| | | *Protaspa* -lineage | *Protaspa* | eukaryvore | testate amoeboflagellate/ amoeba/flagellate (organic/agglutinated) | creeping/gliding on substrate | Howe et al. 2011 |

| | | | | | | | |
|---|---|---|---|---|---|---|---|
| | | Rhogostomidae | | omnivore | testate amoeboflagellate/ amoeba/flagellate (organic/agglutinated) | creeping/gliding on substrate | Dumack et al. 2017 |
| | | | *Capsellina* | omnivore | testate amoeboflagellate/ amoeba/flagellate (organic/agglutinated) | creeping/gliding on substrate | Howe et al. 2011 |
| | | | *Rhogostoma* | omnivore | testate amoeboflagellate/ amoeba/flagellate (organic/agglutinated) | creeping/gliding on substrate | Dumack et al. 2017 |
| | Ventricleftida | | | N/A | testate amoeboflagellate/ amoeba/flagellate (organic/agglutinated) | creeping/gliding on substrate | Howe et al. 2011 |
| | | Ventrifissuridae | | eukaryvore | testate amoeboflagellate/ amoeba/flagellate (organic/agglutinated) | creeping/gliding on substrate | Howe et al. 2011 |
| | | | *Verrucomonas, Ventrifissura* | eukaryvore | testate amoeboflagellate/ amoeba/flagellate (organic/agglutinated) | creeping/gliding on substrate | Howe et al. 2011 |
| | Ebriida | Ebridea | | eukaryvore | testate amoeboflagellate/ amoeba (silica) | freely swimming | Howe et al. 2011 |
| | | | TAGIRI-1-2 | N/A | N/A | N/A | Hoppenrath et al. 2006 |
| | | Ebriacea | | eukaryvore | testate amoeboflagellate/ amoeba (silica) | freely swimming | Howe et al. 2011 |
| | | | *Ebria* | eukaryvore | naked flagellate | freely swimming | Hoppenrath et al. 2006; Howe et al. 2011 |
| | Tectofilosida | | | N/A | N/A | N/A | |
| | | Pseudodifflugiidae | | eukaryvore | testate amoeboflagellate/ amoeba/flagellate (organic/agglutinated) | creeping/gliding on substrate | Howe et al. 2011 |

[1] Also possessing an ephemeral flagellate stage (Phytomyxea) or an amoeboid feeding form and a flagellate locomotive form (some Granofilosea,Viridiraptoridae)

[2] Eulgyphida: There are only two rare exceptions to the silica-scales bearing, *Ovulinata* and *Micropyxidiella,* here neglected.

[3] Euglyphidae nutrition: "herbivore" (=our eukaryvore) in Meisterfeld (2000), algivore (Seppey et al. 2017), *Trinema, Euglypha, Assulina* and *Trachelocorythion* have been fed with bacteria only (Wyzelich et al. 2002), so we consider the family as omnivore.

[4] *Spongomonas* is included in the creeping/gliding category, although it is attached to the substrate.

[5] Cercomonadida are described as bacterivore (Bass et al. 2009) but evidence is accumulating towards myco- and algophagy (references given).

[6] *Proleptomonas* was described as osmotroph (Vickerman et al. 2002), but the ATCC strains 50735 and is fed with bacteria.


Bass D, Howe AT, Mylnikov AP, Vickerman K, Chao EE, Smallbone JE, Snell J, Cabral CJ & Cavalier-Smith T. 2009. Phylogeny and classification of Cercomonadida (Protozoa, Cercozoa): *Cercomonas* , *Eocercomonas* , *Paracercomonas* , and *Cavernomonas* gen. nov. *Protist* , 160:483-521. doi:10.1016/j.protis.2009.01.004.

Brabender M, Keve Kiss Á, Domonell A, Nitsche F & Arndt H. 2012. Phylogenetic and morphological diversity of novel soil cercomonad species with a description of two new genera (*Nucleocercomonas* and *Metabolomonas* ). *Protist* , 163:495-528. doi:10.1016/j.protis.2012.02.002.

Cavalier-Smith T. 1993. Kingdom *Protozoa* and its 18 phyla. *Microbiol Rev* , 57:953-994.

Chantangsi C & Leander BS. 2010. An SSU rDNA barcoding approach to the diversity of marine interstitial cercozoans, including descriptions of four novel genera and nine novel species. *Int J Syst Evol Microbiol* , 60:1962-1977. doi:10.1099/ijs.0.013888-0.

Dumack K, Öztoprak H, Rüger L & Bonkowski M. 2017. Shedding light on the polyphyletic thecate amoeba genus *Plagiophrys* : transition of some of its species to *Rhizaspis* (Tectofilosida, Thecofilosea, Cercozoa) and the establishment of *Sacciforma* gen. nov. and Rhogostomidae fam. nov. (Cryomonadida, Thecofilosea, Cercozoa). *Protist* , 168:92-108. doi:10.1016/j.protis.2016.11.004.

Dumack K, Schuster J, Bass D & Bonkowski M. 2016. A novel lineage of "naked filose amoebae"; *Kraken carinae* gen. nov. sp. nov. (Cercozoa) with a remarkable locomotion by disassembly of its cell body. *Protist* , 167:268-278. doi:10.1016/j.protis.2016.04.002.

Dumack K, Siemensma FJ & Bonkowski M. 2018. Rediscovery of the testate amoeba genus *Penardeugenia* (Thaumatomonadida, Imbricatea). *Protist* , in press

Ekelund F. 1998. Enumeration and abundance of mycophagous protozoa in soil, with special emphasis on heterotrophic flagellates. *Soil Biol Biochem* , 30:1343-1347. doi:10.1016/S0038-0717(97)00266-6.

Flavin M, O'Kelly C, Nerad TA & Wilkinson G. (2000) Cholamonas cyrtodiopsidis gen. n., sp. n. (Cercomonadida), and endo-commensal, mycophagous heterotrophic flagellate with a double kinetid. *Acta Protozoologica* 39:51-60.

Geisen S, Koller R, Hünninghaus M, Dumack K, Urich T & Bonkowski M. 2016. The soil food web revisited: Diverse and widespread mycophagous soil protists. *Soil Biol Biochem* , 94:10-18. doi:10.1016/j.soilbio.2015.11.010.

Hess S & Melkonian M. 2013. The mystery of clade X: *Orciraptor* gen. nov. and *Viridiraptor* gen. nov. are highly specalised, algivorous amoeboflagellates (Glissomonadida, Cercozoa). *Protist* , 164:706747.

Hibberd DJ. 1976. The fine structure of the colonial colorless flagellates *Rhipidodendron splendidum* Stein and *Spongomonas uvella* Stein with special reference to the flagellar apparatus. *J Protozool* , 23:374-385.

Hoppenrath M & Leander BS. 2006. Ebriid Phylogeny and the expansion of the Cercozoa. *Protist* , 157:279-290. doi:10.1016/j.protis.2006.03.002.

Howe AT, Bass D, Scoble JM, Lewis R, Vickerman K, Arndt H & Cavalier-Smith T. 2011. Novel cultured protists identify deep-branching environmental DNA clades of Cercozoa: New genera *Tremula, Micrometopion, Minimassisteria, Nudifila, Peregrinia* . *Protist* , 162:332-372.

Howe AT, Bass D, Vickerman K, Chao EE & Cavalier-Smith T. 2009. Phylogeny, taxonomy, and astounding genetic diversity of Glissomonadida ord. nov., the dominant gliding zooflagellates in soil (Protozoa: Cercozoa). *Protist* , 160:159-189. doi:10.1016/j.protis.2008.11.007.

Meisterfeld R. 2000. Testate amoebae with filopodia. In: *An illustrated guide to the protozoa* (ed. Lee JJ,Leedale G & Bradbury P), Lawrence, Kansas, U.S.A. pp. 1054-1084.

Mylnikov AP, Weber F, Jürgens K & Wylezich C. 2015. *Massisteria marina* has a sister: *Massisteria voersi* sp. nov., a rare species isolated from coastal waters of the Baltic Sea. Eur J Protistol, 51:299-310. doi:10.1016/j.ejop.2015.05.002.

Neuhauser S, Kirchmair M, Bulman SR & Bass D. 2014. Cross-kingdom host shifts of phytomyxid parasites. *BMC Evol Biol* , 14:33. doi:10.1186/1471-2148-14-33.

Seppey CVW, Singer D, Dumack K, Fournier B, Belbahri L, Mitchell EAD & Lara E. 2017. Distribution patterns of soil microbial eukaryotes suggests widespread algivory by phagotrophic protists as an alternative pathway for nutrient cycling. *Soil Biol Biochem* , 112:68-76. doi:10.1016/j.soilbio.2017.05.002.

Shiratori T, Yabuki A & Ishida K-I. 2012. *Esquamula lacrimiformis* , n.g., n. sp., a new member of Thaumatomonads that lacks siliceous scales. *J Euk Microbiol* , 59:527-536. doi:10.1111/j.1550-7408.2012.00635.x.

Strüder-Kypke MC & Hausmann K. 1998. Ultrastructure of the heterotrophic flagellates *Cyathobodo* sp., *Rhipidodendron huxleyi* Kent, 1880, S*pongomonas sacculus* Kent, 1880, and *Spongomonas* sp. *Eur J Protistol* , 34:376-390. doi:10.1016/S0932-4739(98)80007-2.

Thomsen HA, Buck KR, Bolt PA & Garrison DL. 1991. Fine structure and biology of Cryothecomonas gen. nov. (Protista incertae sedis) from the ice biota. *Can J Zool* , 69:1048-1070.

Vickerman K, Le Ray D, Hoef-Emden K & De Jonckheere J. 2002. The soil flagellate *Proleptomonas faecicola* : Cell organisation and phylogeny suggest that the only described free-living trypanosomatid is not a kinetoplastid but has cercomonad affinities. *Protist* , 153:9-24. doi:10.1078/1434-4610-00079.

Vickerman K, Appleton PL, Clarke KJ & Moreira D. 2005. *Aurigamonas solis* n. gen., n. sp., a soil-dwelling predator with unusual helioflagellate organisation and belonging to a novel clade within the Cercozoa. *Protist,* 156:335-354. doi:10.1016/j.protis.2005.07.003.

Wylezich C, Meisterfeld R, Meisterfeld S & Schlegel M. 2002. Phylogenetic analyses of small subunit ribosomal RNA coding regions reveal a monophyletic lineage of euglyphid testate amoebae (order Euglyphida). *J Euk Microbiol* , 49:108.118. doi:10.1111/j.1550-7408.2002.tb00352.x.

**Table S5**. Beta diversity indices calculated for each sampling date.

| | April | May | June | August | October | November |
|---|---|---|---|---|---|---|
| **Total dissimilarity (Bray-Curtis, rarefied data)** | 0.81 | 0.81 | 0.8 | 0.8 | 0.8 | 0.81 |
| **Total dissimilarity (Bray-Curtis, relative data)** | 0.81 | 0.81 | 0.79 | 0.79 | 0.79 | 0.8 |
| **Total dissimilarity (Sorensen, presence-absence)** | 0.53 | 0.52 | 0.52 | 0.45 | 0.46 | 0.54 |
| **Turnover (Sorensen, presence-absence)** | 0.36 | 0.38 | 0.33 | 0.34 | 0.34 | 0.43 |
| **Nestedness (Sorensen, presence-absence)** | 0.17 | 0.15 | 0.19 | 0.1 | 0.13 | 0.12 |

**Table S6**. Linear mixed models showing the effects of the environmental predictors on the most abundant 12 cercozoan families (green=bacterivore; blue=plant parasite, brown=omnivore, black=unknown), the morphotype, nutrition and locomotion modes. We give: a) the spatial correlation structure best correcting the starting model according to the AIC; b) the number of models within two AICc units (after model dredging); c) the number of predictors included in all models extracted in a); d) the remaining, highly significant predictors after fitting a model with just the consensus predictors in b), and their effect type (positive or negative); e) their significance level (p values: *<0.05, **<0.01, ***<0.001).

| Family | Correlation Structure | #Models within 2 AICc units | # Consensus predictors | Highly significant (effect type +/-) | Significance level |
|---|---|---|---|---|---|
| **Sandonidae** | NA | 2 | 4 | Clay (-)<br>pH (+)<br>Soil Moisture (+) | ***<br>***<br>*** |
| **Paracercomonadidae** | NA | 6 | 5 | Archaeal 16S (-)<br>Clay (+)<br>Root Biomass (+)<br>Soil Moisture (+) | ***<br>**<br>**<br>*** |
| **Cercomonadidae** | NA | 13 | 4 | Clay (+)<br>Soil Moisture (+)<br>Extr. Org. Carbon (-) | **<br>***<br>** |
| **Spongomonadidae** | NA | 22 | 2 | Extr. Org. Carbon (+)<br>Root Biomass (-) | **<br>** |
| **Unclassified Euglyphida** | NA | 3 | 4 | Clay (+)<br>Soil Moisture (-) | ***<br>*** |
| **Euglyphidae** | NA | 2 | 5 | C/N Ratio (+)<br>pH (-)<br>Soil Moisture (-)<br>Soil Organic Carbon (-)<br>Total Nitrogen (-) | ***<br>**<br>***<br>***<br>*** |
| ***Spongospora nasturtii* lineage** | NA | 4 | 4 | Microbial Carbon (+)<br>Microbial Nitrogen (-)<br>pH (+) | **<br>**<br>** |
| **Trinematidae** | NA | 2 | 2 | Soil Moisture (-) | *** |
| **Rhogostomidae** | Rational | 17 | 1 | Soil Moisture (-) | *** |
| **Allapsidae** | Spherical | 3 | 3 | Clay (+) | ** |
| ***Polymyxa* lineage** | Spherical | 16 | 2 | Bacterial cell counts (-) | ** |
| **Thaumatomonadidae** | Spherical | 6 | 6 | Soil Moisture (+) | *** |
| **Morphotype** | | | | | |
| **Naked flagellate** | NA | 4 | 3 | Clay (-)<br>pH (+)<br>Soil moisture (+) | ***<br>**<br>** |
| **Naked amoeboflagellate** | NA | 4 | 3 | Clay (+)<br>Soil moisture (+) | ***<br>*** |
| **Flagellate / intracellular parasite** | NA | 5 | 2 | Bacterial cell counts (-)<br>Total N (+) | **<br>** |
| **Naked amoeba** | NA | 23 | 2 | N microbial biomass (-) | ** |
| **Testate amoeba/ amoeboflagellate/flagellate (test organic, agglutinated or siliceous)** | NA | 21 | 2 | Soil moisture (-) | *** |
| **Nutrition mode** | | | | | |
| **Bacterivore**[1] | Spherical | NA | NA | Bacterial cell counts (+) | * |
| **Omnivore** | NA | 11 | 2 | Soil moisture (-) | ** |

| | | | | | |
|---|---|---|---|---|---|
| **Plant parasite** | NA | 2 | 2 | Bacterial cell counts (-) | ** |
| | | | | C microbial biomass (+) | ** |
| **Parasite (not plant)** | Gaussian | 4 | 3 | Total N (+) | ** |
| **Eukaryvore** | NA | 19 | 2 | Bacterial cell counts (+) | *** |
| **Locomotion mode** | | | | | |
| **Creeping/gliding on substrate** | NA | 3 | 3 | Bacterial cell counts (+) | ** |
| | | | | Total_N (-) | *** |
| **Non-motile endoparasite** | NA | 5 | 2 | Bacterial cell counts (-) | ** |
| | | | | Total_N (+) | ** |
| **Freely swimming** | NA | 1 | 2 | Bulk density (-) | ** |

[1] *The basic model for bacterivores included only one significant predictor (bacterial cell counts). Accordingly, a variable selection did not lead to model convergence.*
